# Recurrent Auto-Encoding Drift Diffusion Model

**DOI:** 10.1101/220517

**Authors:** Moens Vincent, Zenon Alexandre

## Abstract

The Drift Diffusion Model (DDM) is a popular model of behaviour that accounts for patterns of accuracy and reaction time data. In the Full DDM implementation, parameters are allowed to vary from trial-to-trial, making the model more powerful but also more challenging to fit to behavioural data. Current approaches yield typically poor fitting quality, are computationally expensive and usually require assuming constant threshold parameter across trials. Moreover, in most versions of the DDM, the sequence of participants’ choices is considered independent and identically distributed *(i.i.d*.), a condition often violated in real data.

Our contribution to the field is threefold: first, we introduce Variational Bayes as a method to fit the full DDM. Second, we relax the *i.i.d*. assumption, and propose a data-driven algorithm based on a Recurrent Auto-Encoder (RAE-DDM), that estimates the local posterior probability of the DDM parameters at each trial based on the sequence of parameters and data preceding the current data point. Finally, we extend this algorithm to illustrate that the RAE-DDM provides an accurate modelling framework for regression analysis. An important result of the approach we propose is that inference at the trial level can be achieved efficiently for each and every parameter of the DDM, threshold included. This data-driven approach is highly generic and self-contained, in the sense that no external input (e.g. regressors or physiological measure) is necessary to fit the data. Using simulations, we show that this method outperforms *i.i.d*.-based approaches (either Markov Chain Monte Carlo or *i.i.d*.-VB) without making any assumption about the nature of the between-trial correlation of the parameters.

## 1. Introduction

Sequential sampling models (1) are models of behaviour that assume that decisions are the result of a noisy accumulation of evidence. The most famous of these models, called the Drift Diffusion Model (DDM), was introduced in the neuroscience field by Roger Ratcliff (2) as a tool for studying animal decision making and has led to an important amount of publications during the past decades.

The DDM is based on a Wiener process, where the evidence in favour of one of several (typically two) actions is represented by a particle moving towards a boundary with a certain drift rate and subject to a given noise. The simplest implementation of the DDM assumes the generative parameters of the model to be fixed across the different trials throughout the experiment. However, this simple version fails to account for some specific patterns of reaction times (RT), which led to subsequent improvements (3; 2; 4; 5; 6; 7) in which parameters are free to vary from trial to trial according to some assumed distribution, giving birth to what is known as the Ratcliff Diffusion Model (8), Extended DDM (9) or Full DDM (10).

While the Full DDM is more realistic (see (11) for a review and discussion), two main drawbacks of this method can be individuated: first, it is more computationally expensive to fit than its simpler counterpart by several orders of magnitudes, as a marginal probability density has to be estimated (or approximated) for each and every trial. So far, several methods used to fit the Full DDM to behavioural datasets have relied on Numerical Quadrature (NQ) (12; 13; 7; 14; 15) or Markov Chain Monte Carlo (16). NQ is computationally expensive, which precludes its use for large problems. While Markov Chain Monte Carlo (MCMC) (17) can be cheaper, it still requires an optimization time and memory storage that is proportional to the size of the problem: this precludes its use on very large datasets, which are encountered more and more frequently in cognitive neuroscience (18; 19; 20).

The second important drawback of the Full DDM regards its identifiability. Typically, only trial-to-trial variability in three (12; 7) or less (5; 21) of the four parameters of the DDM are assessed (drift, start-point and non-decision time, see below). Variability of the fourth parameter, the threshold, is usually regarded as non-identifiable, because its trial-to-trial variability would be equivalent to the variability in the start point position (7; 22). This can be understood intuitively by observing that an increased variability for both these parameters leads to the same behavioural pattern, namely more frequent and faster errors wrt accurate responses. In this perspective, (23) suggest that the choice of the prior distribution shape of the start point (or threshold) determines whether the generative capacity of the model *for a given dataset*. However, the rationale for discarding threshold variability is mainly empirical and lack precise mathematical justification (7): indeed, the lack of identifiability pointed by (23) generalizes to every DDM parameter. We suggest that, even though several models can predict accurately the data at hand, the will differ in their capacity to be generalized to other data and in their model evidence value. Cross-validation and model evidence assessment will therefore be crucial to select model features that should be incorporated in the model. Here, we show that variability in threshold is instead identifiable together with the rest of DDM parameters, especially when the values of the threshold and start point at a specific trial can be predicted from past behavioural observations (time-dependency assumption) and when these values covary with other parameters or physiological measures (covariance assumption). These two restrictions are likely to be observed in practice, as we will see later. In these two cases, one can take advantage of the fact that the values of other variables (latent or observed) constrain the fit of the threshold variability. Although we will focus in the present paper on the first assumption, the approach we propose can model these two phenomena.

Nevertheless, even when the threshold is considered as fixed, identifiability of the Full DDM remains an issue (24). It is usually considered that one can expect the fit of the mean of the DDM parameters to be accurate (7), but not their variance (i.e. trial-to-trial variability) (12). In short, large datasets (e.g. > 100 trials per subject) take unreasonable amount of time to be optimized with these models, for a very limited quality of fit, which worsens dramatically with smaller datasets.

On top of this, another issue with many implementations of the DDM lies in the core assumption of the method used to fit the parameters to the data. Maximum Likelihood Estimate (MLE; i.e. finding the most likely value that the parameters would endorse in order to generate the data at hand) was traditionally used (12; 25; 26; 13; 27; 28; 7; 14), until Bayesian attempts were made (29; 30; 15). Crucially, the validity of the MLE approach is based on some asymptotic properties of the summary statistics of a given dataset when the amount of data tends to infinity (31). However, as we have seen, the computational cost of the numerical methods commonly used to optimize the Full DDM prevents us from fitting large datasets. Hence, we face a dilemma: a large data set will prevent us from computing the ML using classical NQ algorithms, and a small dataset set will violate the assumptions of the MLE. Usage of MCMC only partly solves this problem, as its computational cost is still a function of the size of the dataset, and since it still relies on a strong *i.i.d*. assumption about the generative process.

The core idea of this paper is that trial-to-trial dependency of the parameter values and behavioural results can, and should be used to help estimate the posterior probability of the latent variables. This assertion is mainly theoretically justified by the observation that actions belonging to sequences can not be considered as independent (see for instance (32; 33; 34; 35; 36; 37; 38)). Let us for instance consider the toy example where a participant has been asked to press 9 times the same button (e.g. right), and we are performing inference about the DDM parameters at the 10th button press. From a Bayesian point of view, it is very likely that, at this trial, the subject will choose the same action as before. If the participant is asked to choose, and indeed chooses, the right response, we can expect that the RT at this trial will be, on average, lower than the one at the trials before (?): this is a typical sequential effect. However, if the participant is asked to choose the left response at this trial and responds accurately, it is even more likely that this response will be slower than the previous trials, as the current action violates the previous pattern. Another example is post-error slowing, a well-known psychological phenomenon usually interpreted as a change of speed-accuracy trade-off following an error (39; 40). This might be modeled as a resetting of the threshold to a high value following an error. But an error might also bias the decision in the next trial towards one of the responses, or impair the attention of the subject during the subsequent trials. Our aim will be to take advantage of these cross-correlations in a blind manner, i.e. without making any specific assumption about the precise structure of the trial-to-trial correlation. This will be achieved by capturing the most relevant trial-to-trial cross-dependencies with an assumption-free dimensionality reduction algorithm.

To do this, we first show that Approximate Bayesian inference techniques can efficiently uncover the parameter sequence: the algorithm we propose is a data-driven, nearly assumption-free model that relaxes many classical assumptions, such as the parametric form of the parameter distributions (14) at the within and between subject levels. Also, it can prevent overfitting by the use of dimensionality reduction and it is scalable to large datasets. Finally, training of this model can be trivially parallelized and accelerated using GPU.

More specifically, in this paper, we bring two nested solutions to the aforementioned problems: first, we show that the Full DDM finds a natural formulation in Bayesian Statistics (Section 2.2 and Section 2.3), and that we can use this formulation to solve efficiently this problem using recent advances in Approximate inference, mostly with Variational Inference techniques (Section 2.4). We then present a computationally efficient Variational Bayes model (*i.i.d.-*VB) to compute the marginal probability density of the Full DDM. This algorithm has the advantage of being scalable, its computational cost and memory usage being amortized wrt. the size of the problem.

Next, we show how, when the *i.i.d*. assumption is relaxed, this basic algorithm can be extended to account for trial-to-trial dependency in a data-driven manner using a Recurrent Auto-Encoder (RAE-DDM) (Section 2.6). This approach is assumption-free, in that it does not require the user to select a specific prior distribution for the parameters nor a precise form of trial-to-trial cross-correlation.

We show in the results (Section 3) that our RAE approach outperforms classical approaches (*i.i.d.-*VB and Hamiltonian Monte Carlo, HMC) in terms of model evidence and fit of the data at the trial, subject and population level. An important and compelling result we provide with the RAE-DDM consists in showing that the trial-to-trial variability of the threshold parameter is possible to estimate together with the other parameters of the DDM.

In Section 4, we discuss our model in detail, highlight situations where it might prove useful, see its limitations and introduce potential improvements.

### 1.1. Related Work

Many routines have been proposed to optimize the DDM. Ratcliff and Tuerlinckx (12), and others (41; 42; 10) have used a NQ algorithm with a numerical estimate of the gradient to optimize the likelihood function at the subject level. The computational cost of this method is extremely high, justifying the development of less flexible, but simpler methods (43; 44; 8; 45). Vandekerckhove et al. (30) used a Markov Chain Monte Carlo (MCMC) technique to estimate each trial distribution in a fully Bayesian setting. This is the approach adopted in several Bayesian statistical softwares such as JAGS (46), WinBUGS (30), or Stan (47), which has broaden the use of this technique for the optimization of the Full DDM in many recent publications (e.g. (48; 49; 50; 51; 52)). HDDM is a Python toolbox written by Wiecki and colleagues (15), aimed at fitting the DDM using MCMC. Peculiarly, this toolbox still relies on NQ to compute estimates at the lower level, rather than estimating each local posterior of the DDM parameters. Our aim will be to alleviate the computational burden of the integration process without compromising on accuracy and while keeping the entire original structure of the DDM.

The algorithm we propose differs in many ways from the work of Vandekerckhove and colleagues (30): first, the RAE-DDM is scalable, such that we can make inference about the generative model of large (or very large) datasets with limited cost, as the gradient of the lower bound to the log-model evidence is estimated stochastically. Our use of Amortized Variational Inference (53) allows us to control the convergence of the algorithm using a validation dataset, which is challenging with MCMC techniques. Finally, the RAE-DDM, besides its ability to account for trial-by-trial variability in the threshold values, can also provide a more precise estimate of the posterior distribution of local parameters, as it conditions this posterior on more information than just the RT, the choice and the prior by adding the previous trials as regressors for the value of the prior and local variable. None of the approaches presented so far took advantage of this cross-dependency between the values of latent parameters, choices and reaction times from trial to trial. We are not, however, the first to propose a form of time-dependant estimate of the behavioural parameters: (33; 34; 35; 54) have for instance proposed simpler, informative models to uncover precise psychological heuristics that were always fit using Maximum-Likelihood methods. The assumption-free nature of our model, traded with its latent parameter sparsity, makes it a compelling tool to give a precise and generalizable estimate of the expected value of the DDM parameters at each trial as well as the uncertainty of this estimate.

Also, (55; 21) proposed a model (the Neural Diffusion Model) where the neural data and behavioural parameters are governed by a common distribution that ties together the two kind of measures. Because the covariance of the two types of parameters is inferred, the fit can indirectly take into account the time evolution of the DDM parameters through the observed evolution of the neuro-physiological measure. Notably, this approach requires a large dataset (56) and inference is achieved with MCMC method. Also, it is still based upon a highly informative model, where the user has to set (and possibly test) precisely the form of the model that couples the parameters. Finally, this model is complex and has a large number of parameters, making overfitting an issue.

Our approach, on the contrary, takes full advantage of the dimensionality reduction property of RAE to set the number of parameters of the model to a minimum. It also contrasts with the neural diffusion model by the fact that it is self-contained, meaning that no external input is needed, although it can easily be included in the model. It also relaxes greatly the need to set a precise form to the model that governs the data: as we will see, the use of non-linear mapping between the latent trial-specific prior and its DDM-mapped value flexibly models a large variety of prior shapes with a limited and controllable risk of overfitting.

Our approach to Bayesian inference relies also on a large body of previous work. The Wake-Sleep algorithm (57) developed almost twenty years ago by Peter Dayan constitutes a cornerstone in the history of Bayesian inference. This algorithm is a two-side recognition-generative model, which works in two phases: during the wake phase, the algorithm learns from the data the generative model by minimizing the error between the predictions and the observed data. During the sleep phase, the generative model is held fixed and is used to generate fake data that are used to train the recognition model. Built on the very same idea, Variational Auto-Encoders (VAE) (58) and its recurrent form (59; 60) have quickly become standard and popular tools to model data in many areas (61). Advances in this area go fast, and giving an accurate account of all of them is beyond the scope of this paper. The present paper builds on recent developments of Recurrent Auto-Encoders that have been concomitantly achieved by several groups (e.g. (62; 63)). In short, the first challenge with these techniques regards determination of the best approximate posterior shape for the latent variable **z***_j,n_* (64; 65; 66). An efficient solution to improve the performance of a VAE is to replace the Variational estimator by an Importance Weighted (IW) (67) estimator of the posterior probability. This simple trick can greatly improve the robustness and convergence of the model. We will show in what follows how this technique adapts to the problem of the DDM. Another issue consists in regularizing the parameters of the recognition and generative models: Dropout (68) has been shown to efficiently prevent overfitting with similar models (69), motivating our choice of relying on variational Dropout to perform inference about the value of the network weights (70).

A final remark concerns identifiability of the model we propose, and of the Full DDM in general. The impact of the shape of the prior used for inference has led to vivid debates in the DDM community. Some authors have claimed that a prior probability of an arbitrary complexity can always be found in order to reflect perfectly the data, implying that the choice of the prior completely determines the capability of the DDM to predict data (23). For instance, following these authors’ logic, also disputed by (71), a *J*^th^ component linear mixture model prior (where *J* is the number of trials of the dataset) over the sole parameter ξ can reflect perfectly the dataset, even if the noise parameter is set to 0 and other parameters are kept fixed. However, this overfitting issue applies to any hierarchical model where local variables are modeled: as a matter of fact, there should always exist an MLE estimate of any model that would predict perfectly the data at hand with little or no generalization capability. Indeed, two specific – and widely used – solutions exist to deal with overfitting issues. The first approach consists in using Bayesian Model Comparison (72; 73), in which the model evidence (i.e. probability of the data given the model marginalized over the space of parameters) of models of different complexity is compared. The second approach consists in assessing the model predictive capacity on a *test* dataset, separate from the *training* dataset that is used for fitting the model. In the above example, cross-validation of this ad-hoc model will obviously fail to show that the model is able to provide accurate insights about unseen data. Moreover, we propose another solution to this identifiability problem, which is that the values taken by the DDM parameters at each trial is only random to somne extend, and can, at least partially be predicted based on the values that these parameters (and related data or regressors) took in the past trials. The approach we propose can achieve these three goals efficiently, as the model is scalabe (i.e. it is able to generalize its predictions to unseen data), provides a lower-bound to the log-model evidence, that can be used for Bayesian Model Comparison and can account for autocorrelation of the parameters and data in a blind manner.

Furthermore, the method we propose alleviates the need to specify a precise prior probability distribution, but, by the use of dimensionality reduction, it ensures that the trial-wise latent variable **z***_j_,_n_* captures a maximum of the variability, thus allowing us to keep a highly generalizable model structure with little or no cost caused by the specific prior specifications.

Finally, the capacity of our approach to estimate the threshold variability is a considerable benefit, which has never been considered before. Other authors (e.g. (40; 49)) have used a regression model to estimate the threshold value at the trial level using a measured neurophysiological signal or experimental condition. The RAE-DDM does not preclude the inclusion of these parameters in the model, but would most likely provide a better fit because it also could account for the trial-to-trial dependency in the parameter values that might not be captured by the external regressor.

## 2. Methods

### 2.1. Notation

It is convenient to first define the notation we will use. In the following, we will refer to the trial index as *j* ϵ 1: *J* and the subject index as *n* ϵ 1: *N*. The trial-specific configuration of the five DDM parameters, namely the threshold *a*, the drift-rate ξ, the relative start point *ω* and the non-decision time τ will be grouped under a single vector *ω_j,n_* (or, if applicable *ω_n_* if it is assumed that 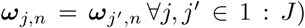. The subject-specific prior distribution parameters of *ω_j,n_* will be generically grouped under the parameter vector *θ_n_*. As we will see in 2.4, we will also define an approximate posterior distribution over *ω_j_,_n_* with parameters *δ_j_,_n_*. In the *i.i.d*.-VB implementation we propose, these two sets of parameters will have a specific form 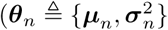 and 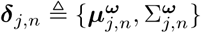). For the purpose of sparsity of the notation and/or to keep the discourse as general as it may, we will use the generic indicators more often than their implementation-specific counterparts.

The behavioural observations are grouped under 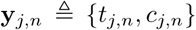, where *t_j_,_n_* and *C_j_,_n_* are the corresponding Reaction time (RT) and choice, respectively. Note that, for the purpose of generality, we distinguish the model boundary *b* and the choice *c*, although we will freely use *p*(*t, c*) and *p*(*t, b*) interchangeably following the context. In general, capital letters will represent the set of all the corresponding parameters or random variables (i.e. 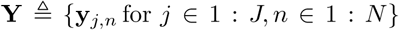,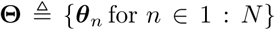 etc.). The *k*^th^ draw of a sample will be indicated by a tilted symbol, such as 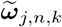, and the true (and obviously unknown) value of a parameter will be indicated by 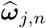.

Finally, if a parameter is deterministically determined by some other variables, such as *δ_j_,_n_* = *f* (**y***_j,n_*; *ρ*), we will use interchangeably this notation with the sparser notation *δ_j_,_n_*(*y_j_,_n_; ρ*).

### 2.2. DDM

The Drift-Diffusion model is a member of the sequential sampling model family. The essence of these models is that the experimenter models the decision process as an accumulation of evidence in favour of one of the available responses: the probability of a pair of observations {*c_j_,_n_,t_j_,_n_*} is formulated as the probability that the evidence reached the threshold *b* at time *t*. Importantly, this accumulation of evidence is stochastic: given the same initial state, the same agent will produce a different response time and, possibly, choose a different item. This stochasticity is the main asset of this model (71), since it can explain speed-accuracy trade-offs: indeed, certain parameter settings will produce faster but more entropic response distributions, whereas other will produce slower, more directed responses.

Formally, in the DDM, the evidence accumulation process is assumed to follow a Wiener process (Figure 1):

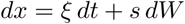

where *dx* is the evidence step, *dt* is the time step and *sdW* a Gaussian noise with a standard deviation *s*. In this specific model, the agent accumulates evidence in favour of an action *c*_1_ with a rate ξ, called the drift rate, meaning that in the case of a classical Two-alternative forced choice task, she will also accumulate evidence in favour of the alternative action *c*_2_ with a rate −ξ. Once the evidence crosses one of the two absorbing boundaries (the boundary *b* positioned at a distance *a* corresponding to the action *c*_1_ or the boundary b at position 0 corresponding to the action *c_2_*), the corresponding action is selected and, after some physiological delay τ, this action is executed. The evidence accumulation can start wherever at *z*_0_ ϵ [0; *a*], although it is more convenient to model the relative start point *w* ϵ [0; 1] where *w* = *z*_0_*/a*. Finally, this 5-parameter model is non-metric: multiplying ξ, *a, z*_0_ and *s* by any positive scalar does not change the behavioural predictions. It is therefore necessary to fix one of them for the model to be used: The accumulation noise *s* = 0.1 is usually used as a metric, but, for the sake of mathematical explicitness and simplicity, we will use the metric *s* = 1 in the following.

**Figure 1:**
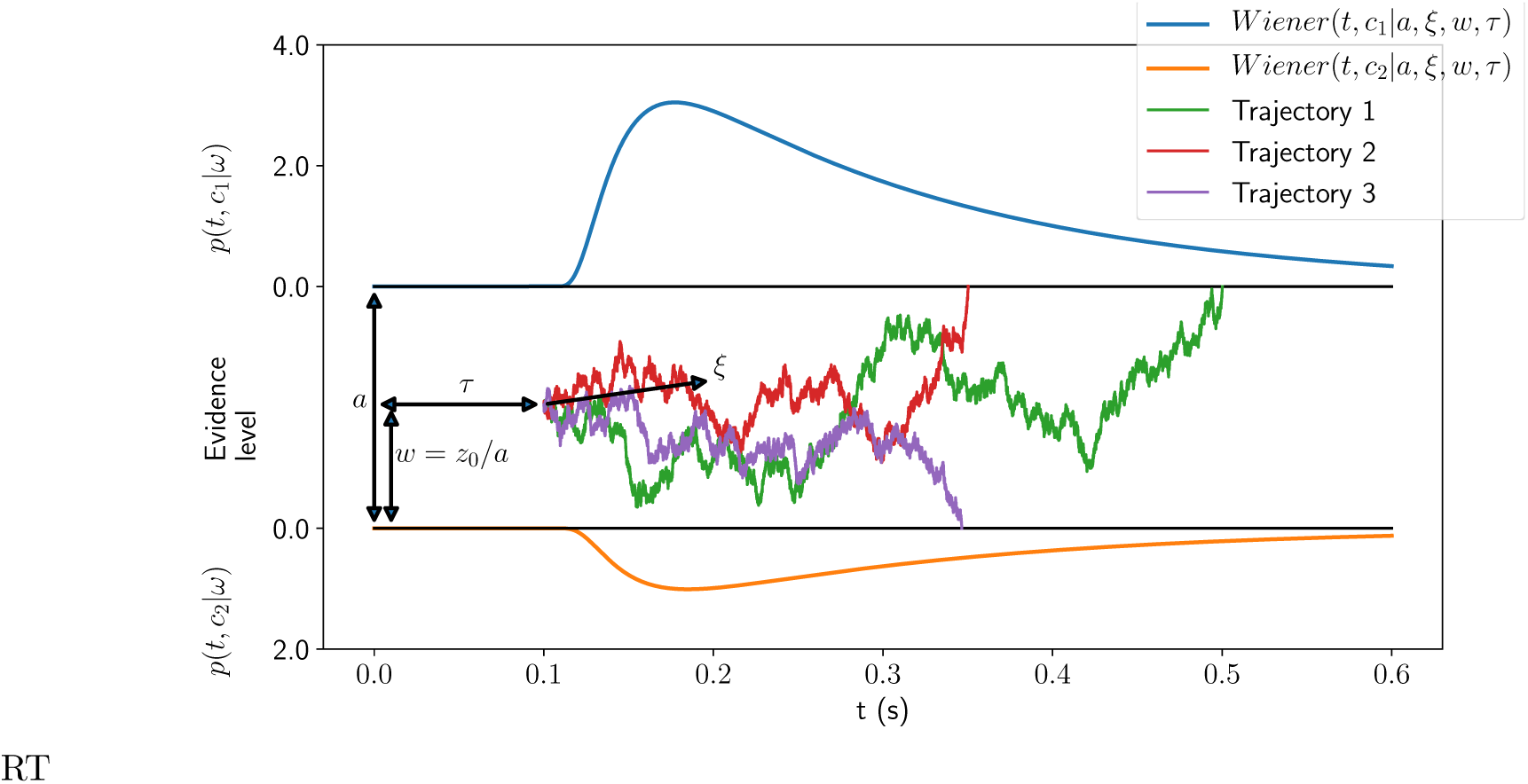
DDM structure. The x axis represents the time scale, the y axis the evidence level. Four parameters (threshold *a*, drift ξ, starting point *w* and non-decision time τ) are represented. At the two boundaries, the first passage probability density function (sometimes called defective density) is represented. Three accumulation trajectories are displayed.

Fortunately, the First-Passage Time density of the Wiener process (*Wiener FPT density*) described before has a closed-form formula. For a Two-alternative forced choice task, the joint probability density of crossing a given boundary *b* (the lower boundary by convention) at a certain point in time *t*, is given by (74):

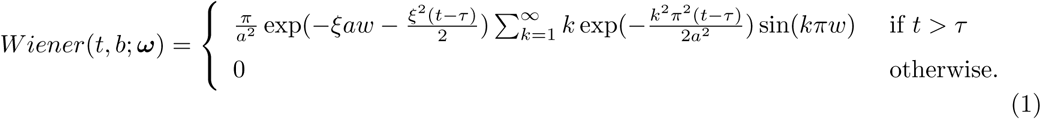

If the absorbing boundary is *b* (i.e. the upper boundary), the above equation can be adjusted by setting *wʹ* = 1 − *w* and ξʹ = −ξ. Together, the sum of the First Passage Time Densities at the two boundaries (sometimes called defective probability densities (6; 21)) forms a valid probability distribution, i.e.:

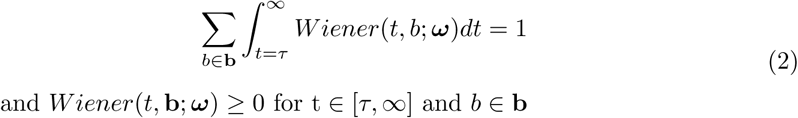

where **b** = {*b*, b}. The function in Equation (1) involves an infinite sum, and must therefore be implemented through approximations. Navarro et al. (75) provided a method to truncate this infinite sum to the appropriate number of terms κ as a function of truncation error. Here, we used their approximation with a numerical precision of 10^−32^ to estimate the parameters value.

The simplest and most intuitive way to fit the DDM to behavioural data is to look for the parameter setting of *ω_n_* that maximizes the likelihood of observing this dataset given the parameters and the model. If tempting, this approach overlooks a crucial detail of the DDM: if the true value of *ω* varies from trial-to-trial, the fit of a single vector 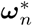 is a biased estimator of 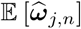, and this bias can have an important impact on the interpretation of the parameter values (5). Formally,

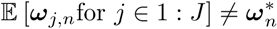

where 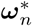 is the value *ω_n_* that maximizes the log-likelihood function.

For instance, a high variability in the value of the starting point *w* with mean .5 will generally produce a dataset that will be better fitted by a point-estimate of *w\** < .5 (5). Accounting for the trial-to-trial variability in *w* can solve this problem. The problem then is to fit a prior ***θ****_n_* that determines the trial-wise distribution of latent parameters *ω_j_,_n_*. In frequentist paradigms, the parameters *ω_j_,_n_* are usually integrated out of the model, and ***θ****_n_* only is fitted. As we will see in 2.3, in a fully Bayesian framework, all the parameters (*ω_j,n_*, ***θ****_n_* and its prior) can be regarded as latent random variables and are estimated accordingly.

Usually, only the subset {ξ, *w*, *τ*} is considered to vary from trial to trial, and the threshold is considered to be fixed for each participant. Although it may raise some concerns in readers familiar with the full DDM, we do no make this assumption here, as it is customary in Bayesian statistics and machine learning to consider that each datapoint is generated by its own set of parameters (see for instance (76)). We show in the results section that the performance of the model we propose still outperforms other implementations, and that the fit of the threshold provides meaningful results.

### 2.3. The Full DDM as an Empirical Bayes problem

The MLE of the Full DDM as formulated by (12) consists in finding the value of the prior 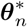 that maximizes the log-likelihood function log *p*(**y***_j,n_*; ***θ****_n_*) while having marginalized out the trial-specific parameters **ω***_j,n_* (and therefore the posterior probability under an unnormalized, unbounded uniform prior), and to consider this value as a consistent estimator of the true value of ***θ****_n_*, namely 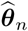. This choice can be justified from the asymptotic normality of the posterior distribution theorem (31; 77): as the dataset grows, and considering that the data are generated from the same process, the posterior distribution approaches a multivariate Gaussian distribution with a mean close to 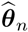 and covariance matrix equal to the inverse of the Observed Fisher Information matrix, i.e. 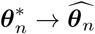 as *NJ → ∞* and 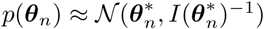, where 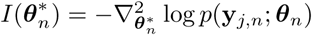. Also, following the Bernstein-von Mises theorem (78; 79; 80), we can see that the choice of the prior becomes irrelevant to the posterior as the amount of data grows, which further justifies the MLE technique for large datasets.

We show in Appendix B that the MLE approach of the Full DDM reduces to what is known as an empirical Bayes estimator of 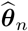 indeed, the value of ***θ****_n_* that maximizes the Wiener density can be directly reformulated as a function of the posterior probabilities of each trial parameter *p*(***ω****_j,n_*|**y***_j,n_*; ***θ****_n_*): this gives the intuition that the problem of fitting the Full DDM could be greatly simplified if one could derive an efficient technique to estimate the set of the trial-wise posterior probability density functions. Figure 2a shows the basic structure of the Full DDM in this perspective.

**Figure 2:**
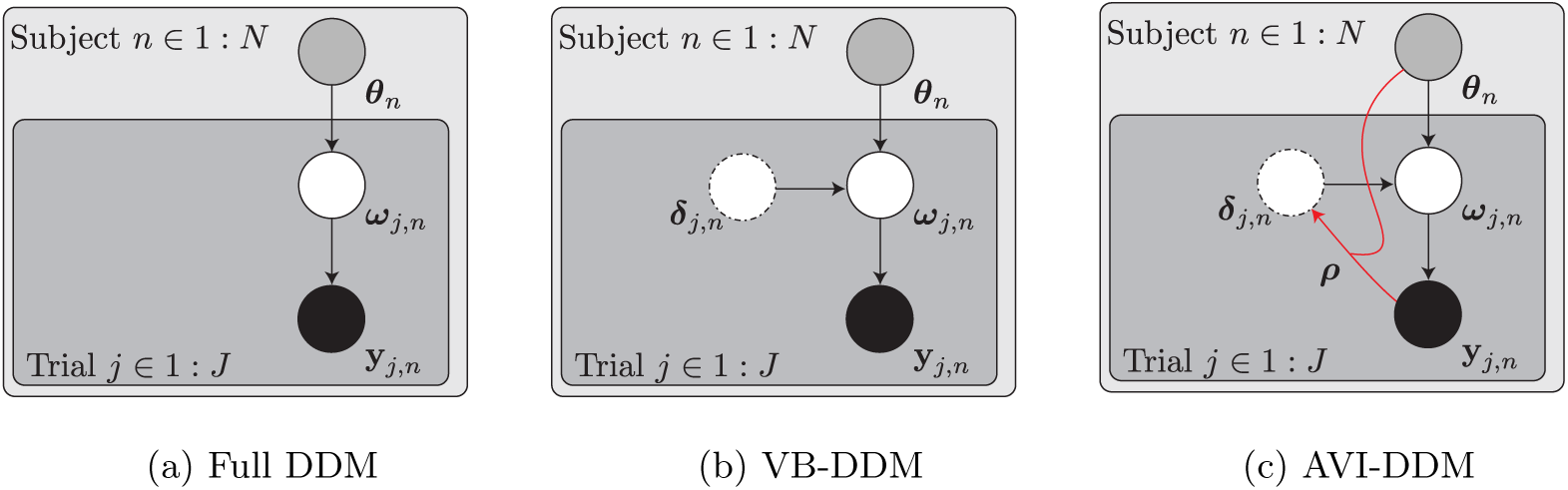
Directed Acyclic Graphical Model (DAG) of the AVI approach to the Maximum-Likelihood (approximate empirical Bayes) Full DDM problem. Black filled circles represent the data (RT and choices), white circles the latent variables, and the greyed circles the subject-specific priors. *a*. Full DDM as a empirical Bayes problem. The aim is to find the value of ***θ****_n_* that maximizes the marginal likelihood *p*(**y***_j,n_*|***θ****_n_*), where ***ω****_j,n_* has been marginalized out. This problem reduces to an empirical Bayes problem. *b*. VB can provide a method to find a lower-bound to the logmarginal probability, by the use of approximate posteriors *q*(***ω****_j,n_w*j,_n_ |*δ_j_,_n_*). *c*. Amortized Variational Inference (AVI) amortizes the cost of inferring the approximate posterior for each latent variable by using an Inference network with parameters *ρ* that takes as input the datapoint and the prior in order to deterministically predict the approximate posterior. Note that, contrary to many similar approaches, here ***θ****_n_* is taken as input to reflect the fact that, with a different prior, the posterior of ***ω****_j,n_* would have a different shape even if the data **y***_j,n_* were identical.

### 2.4. Variational Inference

In this section, we will present Variational Inference as a method to find the posterior of the trial-wise DDM parameters ***ω****_j,n_*. We will first describe the most common EM approach to VB (whose derivation for the DDM application is provided in Appendix B), because it is widely used in neuroscience (81; 82; 83; 84; 85). This EM algorithm consists in looping between the evaluation of each trial-specific posterior probability density function given the current estimate of ***θ****_n_*, before then finding the value of ***θ****_n_* that maximizes the likelihood (or the lower-bound to the log-model evidence in a fully Bayesian setting) given the current posterior estimates. We will expose its weaknesses, and present another, neater method to maximizes the loss function *ℓ*(**y***_j_,_n_*; *θ_n_*) that relies on Stochastic Gradient Variational Bayes (SGVB) and inference networks.

More specifically, the EM algorithm relies on computing the posterior over **Ω***_n_* (the set of all trial-by-trial DDM parameter values), which reads

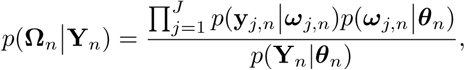

assuming that the posterior probability factorizes over each datapoint. This posterior probability is intractable due to its normalizing factor (the model evidence) *p*(**Y***_n_*|*θ_n_*). A first approach could be to use simulation techniques to approximate this posterior density function, but the size of the problem precludes this choice, since we would need to compute *NJ* independent integrals at each step of the EM algorithm. Instead, we can make use of Variational Bayes techniques (VB) (86; 87; 17; 88) to fit an approximate posterior to the true posterior. In the present case, in which we are interested in the MLE value of the prior, this approach is called Variational EM, or VEM.

In short, VB works as follows: we first define a proxy to the posterior density of an arbitrary shape *q*(***ω****_j,n_*; *δ_j_,_n_*), where *δ_j_,_n_* is a general indicator of the variational parameters that we will optimize in order to match the real posterior as close as possible (see Figure 2b). In order to do so, we can choose to minimize the Kullback-Leibler (KL) divergence between this distribution and the true posterior, since this measure is minimized when the two distributions match exactly:

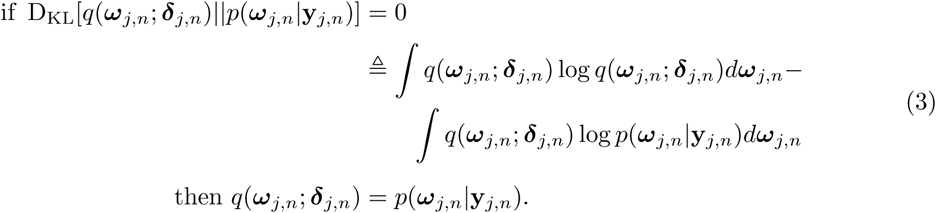

The first term of the KL divergence is the negative differential entropy of *q*(***ω****_j_,_n_;* ***δ****_j_,_n_*):

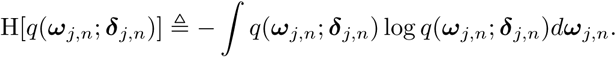

For the second term, we can expand log *p*(***ω****_j_,_n_*|**y***_j_,_n_*) = log *p*(**y***_j_,_n_*, ***ω****_j_,_n_*)*−*log *p*(**y***_j,n_*), and, considering that *∫ q*(***ω****_j_,_n_*) log *p*(**y***_j_,_n_*)*d****ω****_j,n_* = log *p*(**y***_j,n_*), the value of ***δ****_j_,_n_* that minimizes the KL divergence simplifies to:

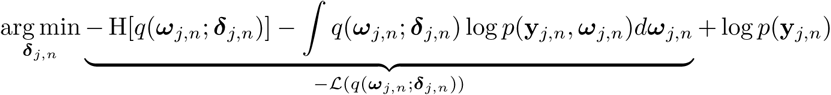

which implies that

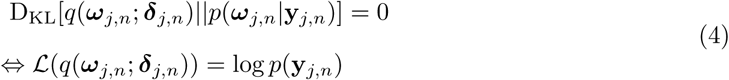

and therefore

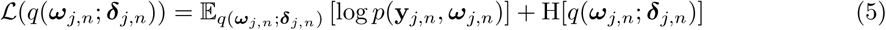

is a lower bound to the log model evidence (or ELBO, for Evidence Lower Bound, also sometimes called negative Free Energy). In essence, Equation (4) tells us that the value of ***δ****_j_,_n_* that minimizes D_KL_[*q*(***ω****_j,n_*; ***δ****_j_,_n_*)||*p*(***ω****_j,n_*|**y***_j,n_*)] is the one that maximizes the ELBO 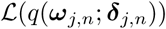. Once this value of ***δ****_j,n_* has been optimized, we can consider that *q*(***ω****_j,n_*; ***δ****_j,n_*) is as close as it could from *p*(***ω****_j,n_*|**y***_j,n_*) given the shape of the distribution we have chosen, the prior probability *p*(***ω****_j,n_*), the data **y***_j,n_* and the likelihood function *p*(**y***_j,n_*|***ω****_j,n_*). Most approaches (89; 90; 91; 92; 93; 94) rely on some form of (possibly natural) gradient ascent method to solve this problem: the gradient of the ELBO wrt. **Δ** = {***δ****_j,n_* for *j* ϵ 1: *J, n* ϵ 1: *N*} is computed, and the optimization proceeds by either optimizing all the variational parameters altogether, or by sequentially optimizing some of them while keeping the others fixed in an EM-like approach. VEM is a special case of the latter, where we are interested in the MLE of the prior.

This VEM method suggested above, where **Δ** and ***θ****_n_* are sequentially optimized, suffers from several issues: the first is that optimizing the approximate posterior for each of the local variables is not trivial, as the ELBO does not have, in general, a closed-form formula (see Appendix B). The second problem is that this approach is highly sensitive to local optima: in other words, the initial draw of **Δ** can frame drastically the final result of the optimization process (17). Another important concern is that the cost of such optimization process is unreasonably high, as one needs to iterate across every datapoint in order to make a single update of the prior, which is not guaranteed to be significant. All these obstacles find an elegant solution with the combined use of SGVB and Inference Networks.

#### 2.4.1. Stochastic Gradient Descent, Reparametrization trick and SGVB

Let us rewrite the optimization problem we need to solve to find the MLE of ***θ****_n_* using VB in a concise form:

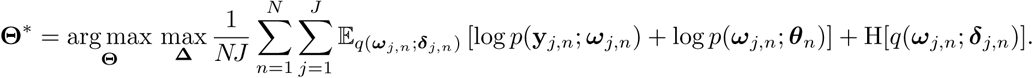

The VEM approach detailed above essentially reduces to maximizing the ELBO wrt. **Θ** and **Δ**. Intuitively, a more economical way to achieve this would be therefore to maximize both parameters at the same time. The gradient of this expression can be expressed exactly as

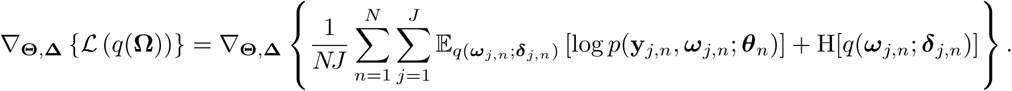

With a little loss of generality, we consider that the entropy of the approximate posterior has a closed-form formula, and that samples can be generated from *q*(***ω****_j_,_n_*; *δ_j_,_n_*) using a differentiable function *g* st. 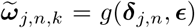 where ϵ ~ *p*(ϵ) for some arbitrary *p*(ϵ). Under these conditions, the above expression has an unbiased, noisy Monte-Carlo estimator given by

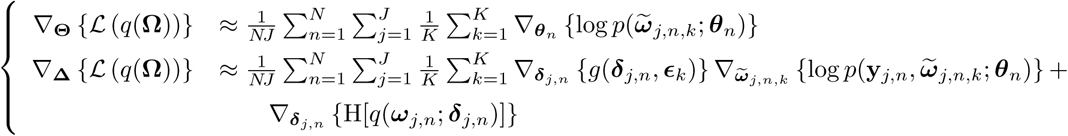

for a number of *K* samples. The form we have given to the (expected) log-likelihood makes it clear that

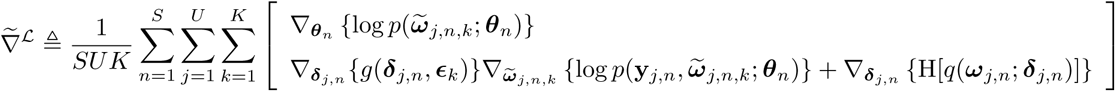
is an unbiased, noisy estimator of 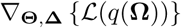, for any *S* ≤ *N* and *U* ≤ *J*.

The use of the auxiliary variable to compute the gradient estimator is sometimes called Infinitesimal Perturbation Analysis estimator (95; 96; 97) or Reparametrized Gradient (98). It has generally a much lower variance than the gradient obtained with the Score function method (98; 93), and, for a large enough mini-batch size *US >>* 0, it can be used with a single sample *K* = 1.

This kind of noisy gradient can be used to find the minimum of a loss function using Stochastic Gradient Descent (SGD) techniques (99), which have numerous advantages. First, as our notation suggests, we can subsample some trials in the whole set in order to maximize the ELBO wrt. both **Θ** and **Δ**. This already amortizes the cost of the optimization, as we do not need to go through the whole dataset to make a step toward the maximum. Second, we do not need to compute the true posterior probability of each of the sampled trials in order to estimate the gradient of the respective variational parameters ***δ****_j,n_*: by sampling from this approximate posterior and computing the gradient of this sample, we can estimate the gradient of the ELBO wrt. ***δ****_j,n_*.

Finally, as for the VEM approach, there exists a risk of ending in a local optimum. This risk is, however, mitigated by the use of SGD: as only part of the data are used at each time step to climb the likelihood curve, local peaks will usually be avoided because they are caused by a small cluster of data that is partially ignored during this step (100).

Many SGD algorithms have been developed (e.g. (101; 102)). As our aim is not to make an extensive review of their respective advantages and disadvantages, we decided to use the Adam optimizer (58) and refer to the corresponding paper for a detailed algorithmic description and implementation.

#### 2.4.2. Inference Network

Flexibility and sparsity are two of the main desired properties of the approximate posteriors q that we want to fit to our dataset: a loss of flexibility will decrease the performance of the model, or equivalently, worsen the unknown KL divergence between the approximate and true posteriors. Sparsity is however the main asset of Variational Inference, and one would wish to find a cheap form of approximate posterior to perform inference in a reasonable amount of time. A solution to reduce the number of variational parameters consists in using a direct, deterministic mapping from the values of the adjacent nodes of factor graph to a given approximate posterior 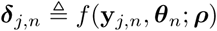 where ***ρ*** is the vector of parameters of the arbitrary function *f* that achieves this mapping. This leads the approximate posterior to have the form *q_ρ_*(**z***_j,n_*|**y***_j,n_*, *θ_n_*) (Figure 2c). Our goal has now become to optimize the mapping function parameters ***ρ*** instead of the local variational posteriors ***δ****_j,n_* which it determines.

This use of an inference network (IN), known as Neural Variational Inference and Learning (53) or amortized variational inference (AVI) (64) has led to numerous applications (93; 103; 64; 70; 104; 65; 105), as it has several advantages. First, from a computational point of view, it alleviates the need of optimizing each of the local approximate posteriors, which can be problematic for large datasets. Instead, a generic function that maps the points to their approximate posterior, is fitted to the problem at hand. Second, this technique allows the model to find a likely posterior for datapoints that might not have been seen yet, or that have been seen many iterations ago. Finally, and following the same logic, algorithms using IN can spare large amount of memory usage by storing only the parameters of the mapping ***ρ*** and not each of the local variational parameters.

Optimizing the ELBO with an IN is straightforward: using the above definition of the approximate posterior, the noisy approximation to 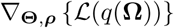 becomes

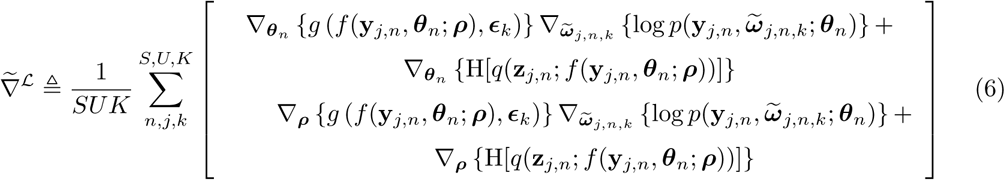
where 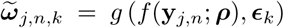. The gradient in Equation (6) can be computed easily using an Automatic Differentiation routine if *g*(·) is differentiable, which excludes for instance random number generation algorithms based on acceptance-rejection algorithms (106; 107), although this limitation can be circumvented (see (108; 109)).

*Approximate Posterior specifications*. We now need to specify a convenient shape for the approxi-mate posterior of the DDM parameters. In Appendix C, we opted for the use of Normalizing Flows (69) as a highly flexible albeit simple distribution shape. We found that, for the DDM, this form of posterior distribution outperformed generally simpler approximate posterior (such as Gaussian distributions) in terms of KL divergence to the log-model evidence.

### 2.5. Preliminary summary

So far, we have presented a method to compute the maximum likelihood of **Θ** using Variational Inference. Through the use of SGVB, we propose an optimization method that has a limited cost for big datasets: indeed, the number of samples per gradient estimation *S* * *U* * *K* depends more on the complexity of the problem than on the size of the dataset. Also, it alleviates the need to compute the marginal likelihood to get a gradient estimate. Finally, it should be robust to the presence of local optima, thanks to the use of AVI, although this cannot be guaranteed. The use of IN scales down the computational burden of the problem, and ensures a certain similarity between the approximate posteriors of each datapoint. Moreover, it makes it possible to optimize the algorithm online, as the same mapping function as the one used for observed datapoints can be used for datapoints that have not been observed yet.

Comparison of Algorithm 1 and Algorithm 2 (see below) highlights the difference between the VEM method and AVI. Note that, in the former, the exact posterior probability from Equation (B.6) has been replaced by its approximation *q* whereas in the latter, the update have been formulated in terms of gradient *ascent* for simplicity of the notation.

Of course, the naive implementation presented in Algorithm 2 does not render the full richness of SGVB: the stopping criterion can, for instance, be refined st. the optimization process is stopped when the performance on a validation dataset starts to decrease (early stopping (17)). This is the method we will present in the Result section 3. Also, the learning rate *η* can be adapted to match an approximation of a preconditioning matrix, which is indeed what the Adam optimizer, mentioned above, does.

**Algorithm 1:**
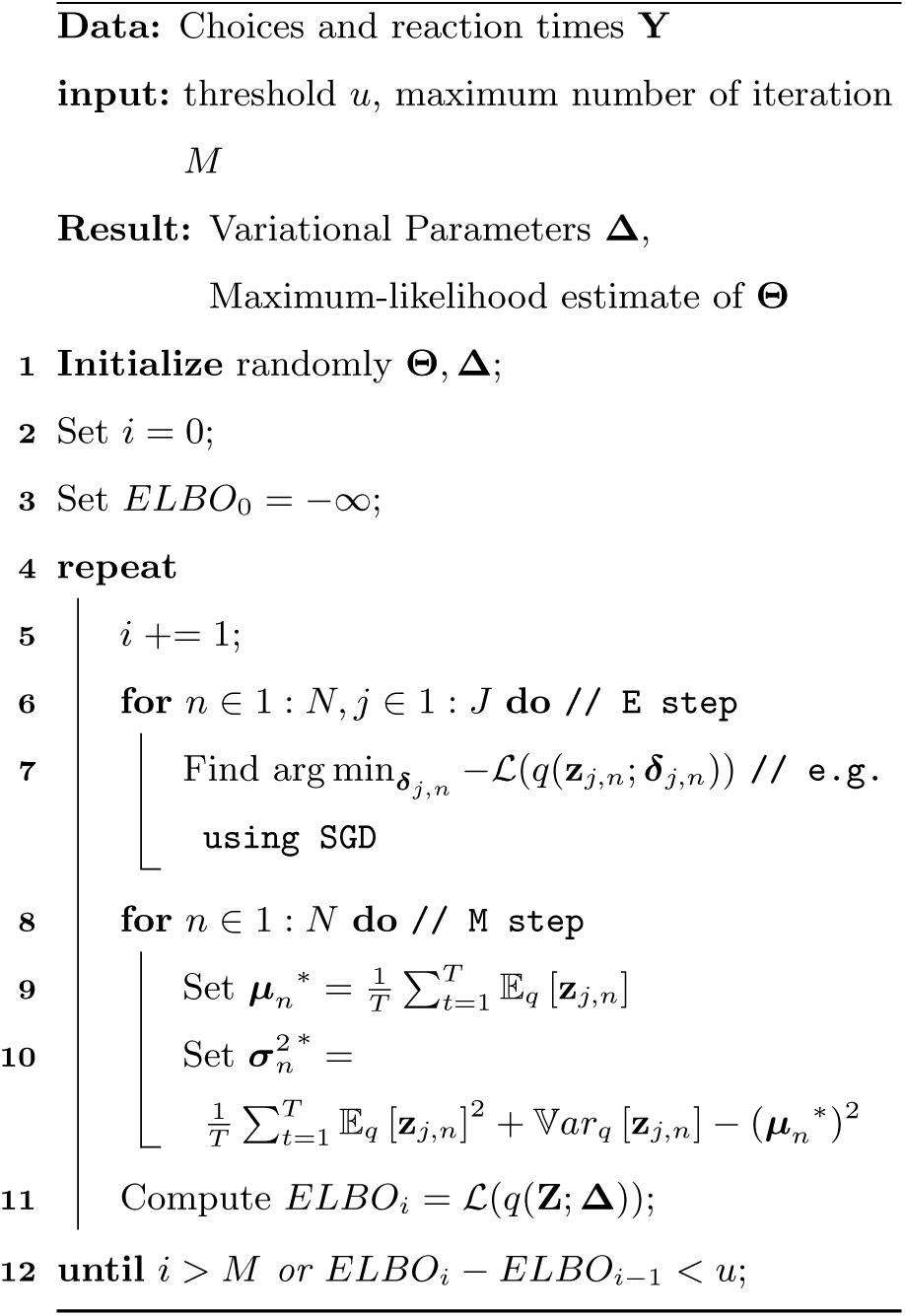
VEM algorithm.

**Algorithm 2:**
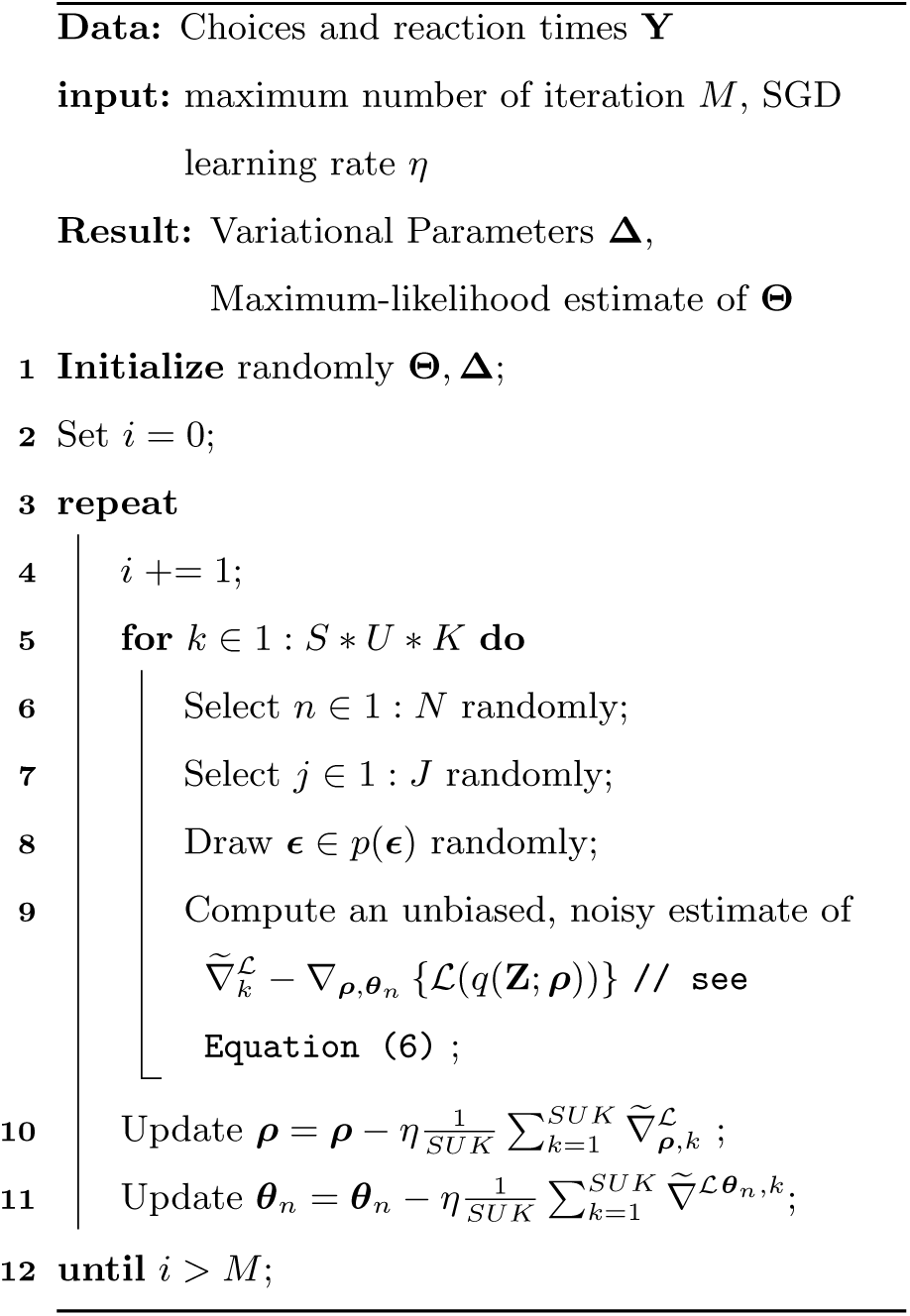
SGVB Amortized Inference

### 2.6. Recurrent Auto-encoder

We now develop the core idea of this paper, which is that one can take advantage of the sequential nature of data acquisition to predict the trial-wise values of the DDM parameters.

The model we will develop is based on Variational Auto-Encoders. Auto-Encoders (AE), and more specifically Variational AE (VAE), are a large family of models (93) that can be seen as two-side algorithms were one side (the *encoder* or *recognition model*) is the equivalent of what we have called an *inference network* so far: it is a (non-)linear mapping with global parameters ***ρ*** that determines the shape of the approximate posterior of a local latent variable **z***_j,n_* of an arbitrary length *d*, based on the data **y***_j,n_*, with a prior *p*(**z***_j_,_n_*). This variable **z***_j,n_* lies between the *encoder* and the *decoder*, which constitutes the *generative model* about which inference is made. Figure 3a sketches the structure of a Variational Auto-encoder.

This generative model maps the parameter **z***_j,n_* onto the parameters of the distribution of **y***_j,n_* (***ω****_j,n_* in our case) thanks to a (non-)linear deterministic mapping ***ω****_j,n_* = *f* (**z***_j,n_*; *ϕ*).

This peculiar model class has the advantage that it allows the user to specify the degree of complexity of the inference process she is trying to achieve by adjusting the hyperparameter *d*. Indeed, VAE ultimately boil down to dimensionality reduction algorithms, where the bottleneck is situated at the level of the latent variable **z***_j,n_*.

Formally, a VAE for a trial indexed *j, n* is defined by the following formula of the ELBO:

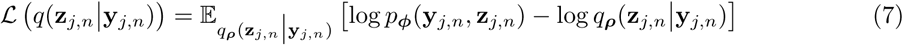

which recalls Equation (5) with the only difference that now an encoder/decoder scheme is used to link the data to the latent variable and the latent variable to the data.

We kept on purpose the subject indices in Equation (7), and the hierarchical treatment of this expression will be introduced shortly after. Let us emphasize in Equation (7) the dependency of the probability distributions on ***ρ*** and *ϕ*: these are the only parameters to fit in order to maximize the ELBO. Ideally, one would like to make inference not only about **z***_j,n_* but also about *ϕ*, as these parameters also belong to the generative model. Often, however, authors treat these parameters in a Maximum-A-Posteriori (MAP) perspective (93), although inference about these parameters is far from being impossible (70; 65). This problem will be treated in 2.6.1.

Chung and colleagues (60) have shown that one can include in this model a specific form of regressor that deterministically brings to the current trial the information about the state of the model at the trial before: this approach, known as Recurrent VAE (RVAE), has been the focus of several recent publications (59; 60) due to its efficiency to trade off accuracy with complexity of the inference process. Under this assumption, the ELBO now looks like:

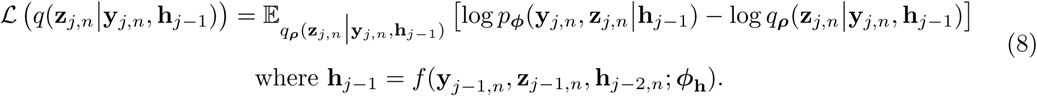

In Equation (8), we have made explicit the fact that the approximate posterior *q* and the likelihood *p*(**y***_j,n_*, **z***_j,n_*) depend on the previous value of **z, y** and **h** through a hidden state **h***_j_*_−i_. As our notation suggests, **h** is also computed using a (non-)linear transformation of these variables, that takes as input a subset of the decoder parameters *ϕ*.

This model specification can be understood as follows: we are interested in retrieving a posterior probability distribution of a variable **z***_j,n_* that acts as regression weights of regressors **h**, which brings to the current trial a selection of information about the trials before (either parameter values, external regressors such as experimental conditions, or behavioural measure, RT and choices included). By specifying the length of **z***_j,n_* to its minimum, we ensure that each element of **z***_j,n_* explains the maximum of the variance of the output parameters of the DDM, ***ω****_j,n_*. Due to the form of the regression we use (i.e. multi-layered perceptron), we can get high flexibility in a space of reduced dimension: very rich patterns can be reflected by such a scheme, even in a low-dimensional space.

Note that we do *not* specify the distribution shape of ***ω****_j,n_*, but the distribution of **z***_j,n_*, which then maps non-linearly to ***ω****_j,n_*. This contrasts with all DDM implementations we are aware of, where the choice of the DDM parameter prior is mostly arbitrary.

Regarding the choice of a prior for **z***_j,n_*, one can assume that it is normally distributed in Equation (7) with a mean 0 and a covariance *I_d_*, which ensures that the components of **z***_j,n_* are somehow independent and, therefore, that each of them captures the maximum variance in the data. However, this choice is not optimal for the model in Equation (8), as one would expect the prior of **z***_j,n_* to vary from trial to trial. We used therefore the recommendation of (60) and used a form of prior 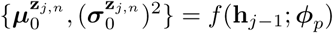 where *ϕ_p_* is again a subset of the generative model regression weights and biases.

The final model specification we need to consider is the way we will capture inter-individual differences. This can be efficiently achieved using a subject-wise auxiliary latent variable **ζ***_n_* of length *d\** with a prior 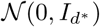 and an approximate posterior *q*(*ζ_n_* |*υ_n_*). This regressor was introduced in the generative model described above to obtain a set of DDM parameters that depended on the three factors **z***_j,n_*, **h***_j−i_* and **ζ***_n_*:

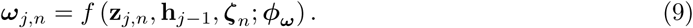

This simple manipulation ensured that inference at the trial level was a compromise between subject-wise behavioural specifics and consistent population-wise behavioural patterns. It goes beyond saying that if some regressor is added to the decoder, it should be added to the encoder too to improve inference about the local variables, which makes us define the following amortized family of approximate posterior over **z***_j,n_*

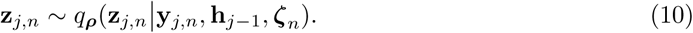

It flows directly from Equation (9) and Equation (10) that, if the dataset has been acquired using various experimental conditions, or if the subject is somehow informed about its performance in the task by the display of a sensory feedback, these regressors can be easily included in the set of the predictors of ***ω***, making this model a compellingly generic tool for inference about the vector ***ω***.

*Training the RAE-DDM*. All the tools we have provided so far can be used in order to train a VAE: the inference network, with possibly the normalizing flow that is presented in Appendix C can be used to find the posterior probability of **z***_j,n_* with an amortized cost. SGVB (by the use of the reparametrization trick) could be used to optimize the ELBO, with an unbiased gradient of the form

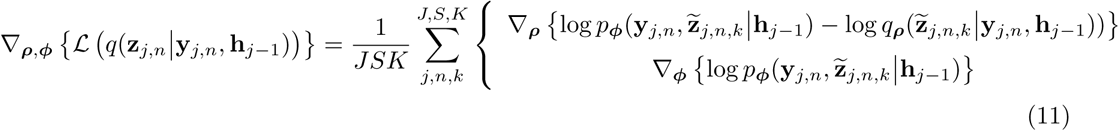

where we have made implicit the fact that **z***_j,n_* was generated according to a IN approach with 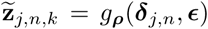. Using this approach, one would generate *K* independent random sequences of 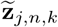 for *j* ∈ 1: *J*, but this approach might suffer from several issues: specifically, the error of each sample will propagate through the whole sequence, and the effect of an early ”bad” sample will impact all the successive trials, which will severely impair the convergence of the SGD algorithm.

**Figure 3:**
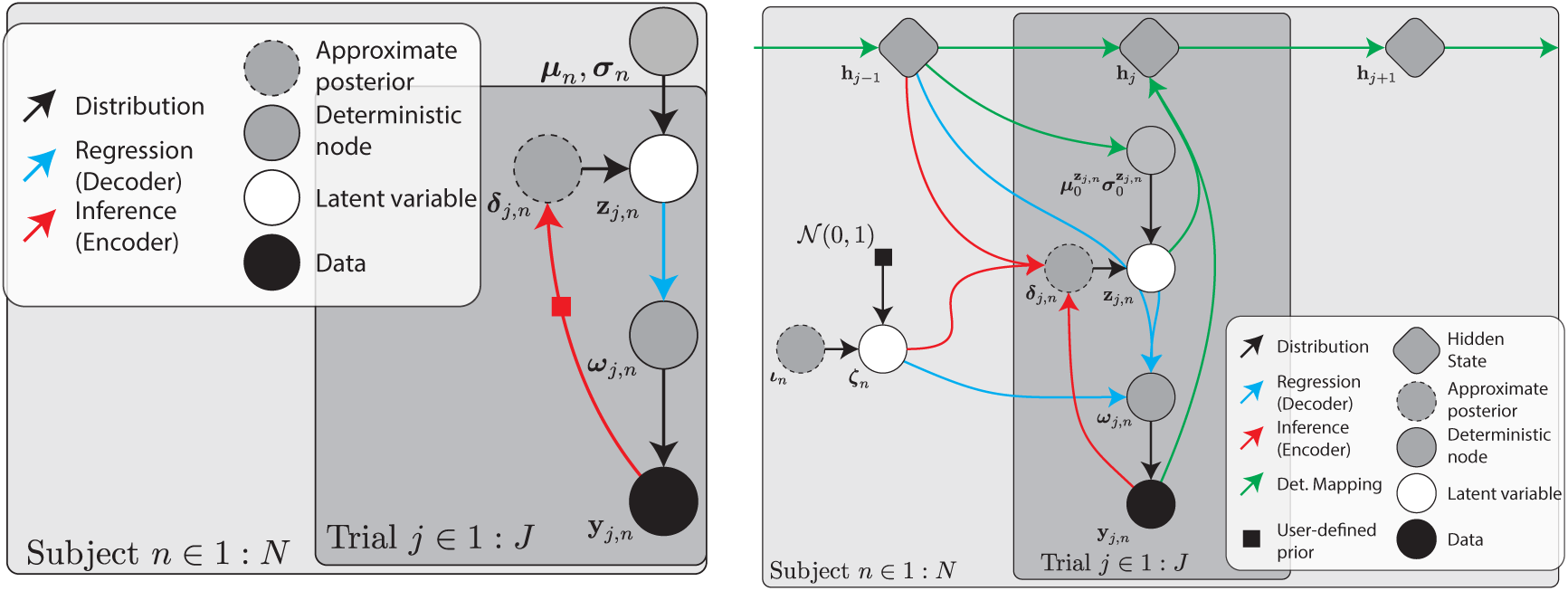
(a) Variational Auto-encoder with Maximal-Likelihood estimate of the top-level hyperprior. Note the difference with 2c, where the latent variable directly conditioned the data **y***_j_,_n_*. (b) Recurrent Auto-Encoding Drift Diffusion Model structure. The whole encoder-decoder process is shown for a trial *j* and a subject *n*. The model is fully connected, in order to account for dependency between the adjacent levels at the recognition and generative level.

Another issue, closely related to the previous one, regards the sampling of the non-decision time *τ*: it is hard to put constraints on the output of the generative regression network, and some samples of *τ* might fall beyond the *t*, where no gradient can be computed.

To solve these problems, a typical solution can be to move the sampling average *inside the logarithm function*, which transforms the ELBO in an Importance Sampling (IS) estimate of the the log-model evidence:

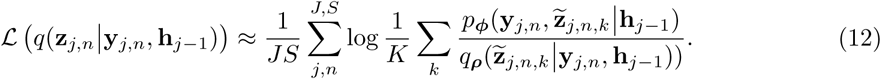

When used without recurrent predictions, this is known as an Importance Weighted AE (IWAE) (67), and when used in the context of sequential data, this algorithm can be viewed a special member of the Particle Filtering family of algorithms which has been named Filtering Variational Objective (FIVO) (62).

This improved estimate of the ELBO has many advantages, that are detailed in (62): it allows us to weight each sequence of data to compute the approximate log-model evidence, and also to drop and resample sequences that fail to reflect the data in a meaningful way based on the other sequences. Moreover, this approach produces samples with a much lower variance than the SGVB approach, which can be simply understood by observing that the FIVO approach takes more samples than SGVB, which is usually taken with a local sample size of *K* = 1. Finally, this approach solves the problem of the bounded space of τ: we can assign a log-likelihood of −∞ (i.e. a weight of 0) to the particles *k* whose τ*_k_* fall beyond the *t*, and provided that at least one the particles does satisfy the conditions for the log-likelihood to be computed, the gradient of the estimator can be computed. However, it might still be the case that the whole set of particles at a given trial produces a sample 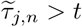. We describe in Appendix C.3 a simple solution to this problem based on Lagrange multipliers.

The complete model, which we name the Recurrent Auto-Encoding Drift Diffusion Model (RAE-DDM) is an application of the algorithm presented in (60; 62) to the problem of DDM fitting, and we refer to these articles for the implementation details. The model is sketched in Figure 3b and the training algorithm is displayed in ??.

**Algorithm 3:**
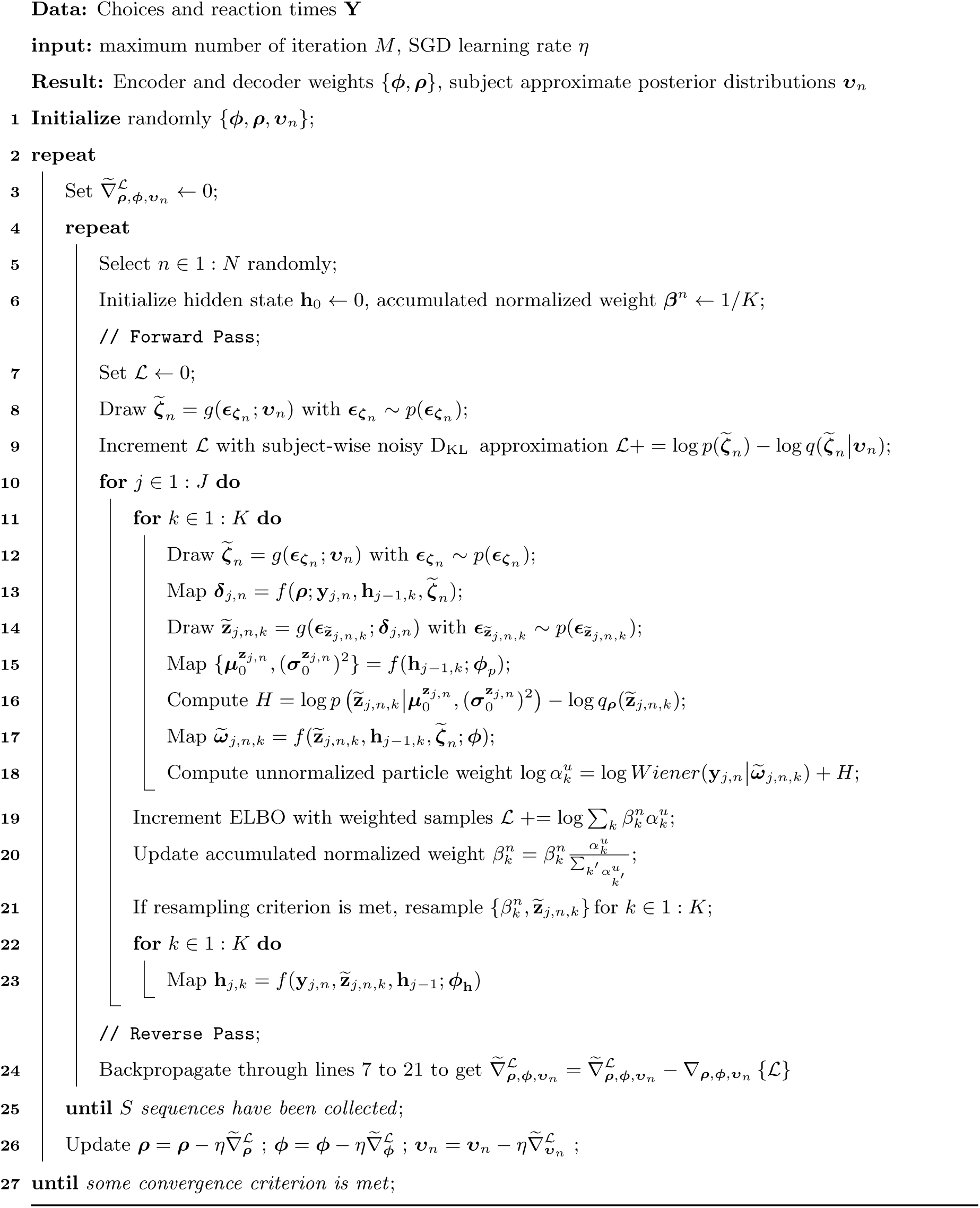
RAE-DDM Training

#### 2.6.1. Variational Dropout

A final consideration regards the overfitting propensity of RNN and, consequently, the potential overfitting of the RAE-DDM. Dropout is a popular technique to prevent overfitting when training a neural network (68), and has been suggested as a necessary requirement for Boltzman-machine like algorithms (69) such as VAE and RAE. It simply consists in hiding part of the network at each iteration, in order to ensure recursivity in the network and maximum relevance of each of the inputs. Variational Dropout (VD) (70; 110) is an improved version of the Dropout method where a specific prior and corresponding variational distribution is assigned to each weight of the network. Using VD, we can ensure that the model takes maximum advantage of each of the inputs of the network, and prevents overfitting. Additionally, the use of VD makes the model fully Bayesian, as we make inference not only about **z***_j,n_* but also about the weights ***ρ*** and *ϕ*. VD implementation of the RAE-DDM performed slightly better than the vanilla version on the long-run, and prevented the optimization to fall into spurious local posterior estimates of the DDM parameters. For each set of weights, including those of the GRU RNN algorithm, and according to (111), we sampled 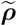 and 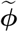 according to their posterior distribution before achieving the whole sequence of forward/backward pass described in Algorithm 3, meaning that we used the same sample for each and every trial of the sequence instead of resampling at each trial.

## 3. Experiment

### 3.1. Data Generation Procedure

To provide a proof of concept of the usefulness of accounting directly for the trial-to-trial dependency of the DDM parameters, we generated a series of datasets with a random Gaussian-Inverse-Gamma prior over the DDM parameters (see Section 3.1), so that the average mean value of each parameter was variable across datasets. The values were chosen with the following heuristic: we ensured that the chosen parameters generated datasets with a good accuracy in the simulated task (see below) with reasonable RT. We always aim at a high trial-to-trial variability of the *ξ* and *w* (lower-bounded at 0.3), and the variability of *τ* and *a* were chosen to be low (bounded at 0). These choices were motivated by the fact that we did not expect the non-decision time to vary more than by a few milliseconds. We also wanted to investigate whether the threshold variability was identifiable for low-to-moderate values of the variance, as this variability is usually assumed to be equal to zero in the DDM literature. The simulated task consisted of a highly generic TAFC task where the subjects needed to choose between a left and a right response to indicate, for instance, the motion direction of dots in a random kinetogram. There was no other manipulation imposed on these datasets. We assumed that the motion direction affected the drift and the drift only. This is a quite common assumption in the DDM literature (7), which is physiologically sensible, as the start point should be set before the instruction is perceived by the agent (112).

**Table 1:**
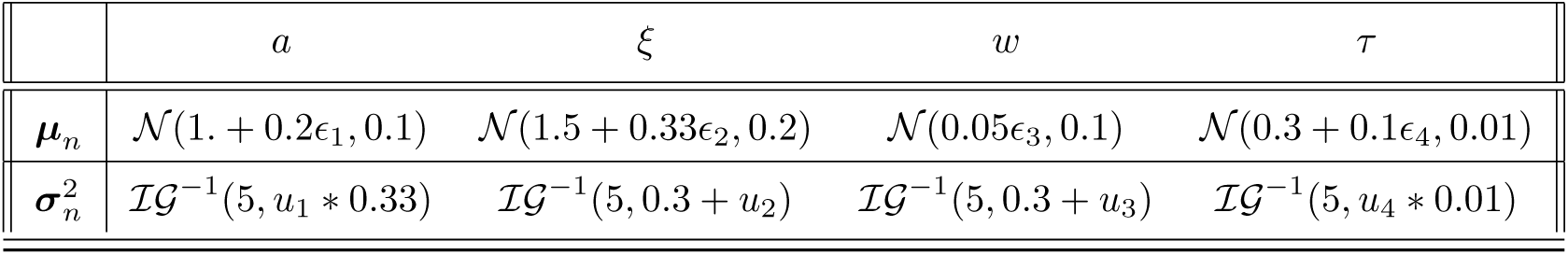
Generative distributions at the population level (i.e. dataset-specific). 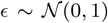 and 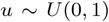 are random noise variables that controlled the distribution of the parameters at the population level. This manipulation ensured heterogeneity in the datasets.

In order to avoid biasing the data generation process by using a generative model that would only be identifiable using a non-*i.i.d*. based approach, the only manipulation we allowed ourselves to do was to permute the parameters, initially generated according to *i.i.d*. normal distributions (Figure 4). This permutation induced specific patterns of cross-correlations between successive parameter values, while ensuring that a fit achieved using the *i.i.d*. assumption was not harmed by the assumption of the data generation procedure.

**Figure 4:**
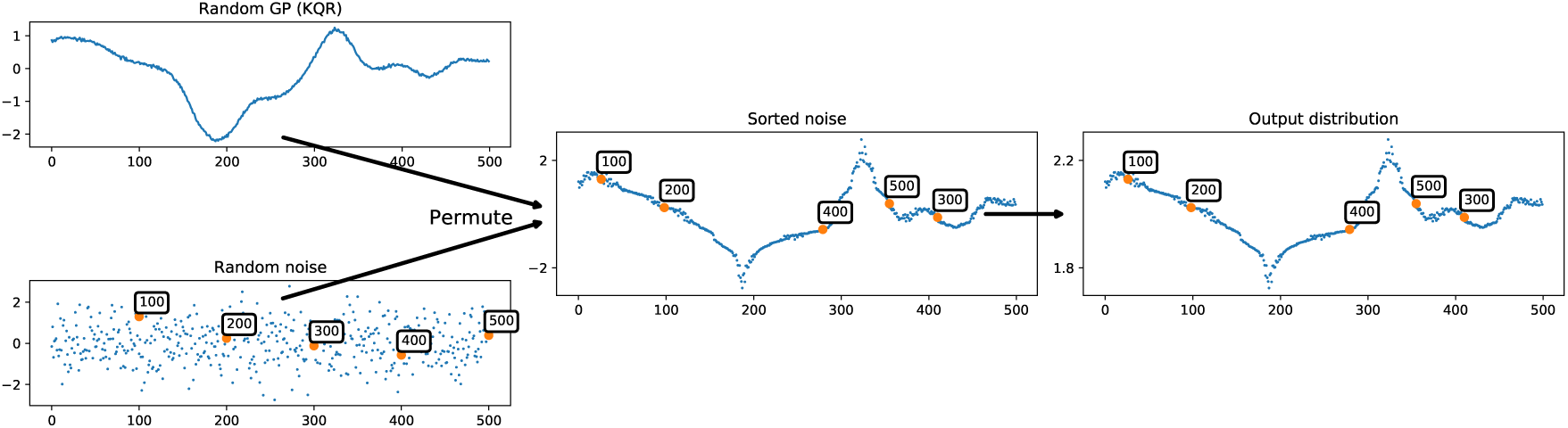
Data generation procedure for a prior distribution 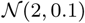. All parameters were generated according to a diagonal univariate Gaussian distribution. For each parameter, the order of the appearance of each sample was permuted to minimize the distance wrt. a random GP trace (here with a Rational Quadratic kernel).

We proceeded in the following way: first, we generated a random matrix of unit-Gaussian distributed random variables 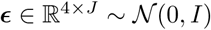. Then, we generated a trace of 4 signals following a Gaussian Process (GP) with a Rational quadratic or a Periodic Kernel (113), parametrized with a population-wise random parameter *a* ~ [0; 1] and with an input vector of equally distant variables u situated between −4 and 4. This simple manipulation ensured that the frequency of the parameter variability was mostly identical in the whole population but different between datasets, while keeping a rich family of traces form across subjects. Next, we permuted the elements of each column of e in order to minimize the Euclidean distance between each of these elements with the correspondent point generated by the GP. Finally, the data were mapped to their desired distribution location by simply applying the following equation:

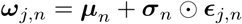

where 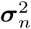 is the Hadamard product operator.

RT and choices were simulated according to (12), where the Wiener process is discretized to an arbitrarily small *δt* (set to 10^−6^ in our case) and approximated as a binary gambler ruin problem (74). An alternative method could consist in using the Euler-Maruyama algorithm (114; 115), but this algorithm did not prove to be of greater accuracy than the former, for a higher computational cost.

A total of 16 datasets of 64 subjects each and 500 trials per subject were generated with the same procedure, and different arbitrary random seeds (from 1 to 16) for reproductibility.

Each dataset was fitted to the RAE-DDM, to a hierarchical *i.i.d*.-VB version of the DDM (see below) and finally to an HMC version of this last model coded in Stan (47). A single chain per dataset was generated, with 1000 iterations for warm-up and 1000 iterations for sampling. Because the threshold was generated with a random noise (as every other parameter), and because it is customary to fit the DDM while discarding the threshold variability, we also fitted the fixed-threshold of the *i.i.d.-*VB -DDM and similarly for the HMC version in order to assess how good the classical threshold steadiness assumption worked. For convenience, these fits will be named “classical” in what follows, by contrast with the “full” versions that included the threshold variability. This leads to a total of five fits per dataset.

### 3.2. Model Specification

*i.i.d.-VB*. For each of the *f* (·) transform in Section 2.6, we used a two-layer perceptron, except for the mapping of **h** for which a Gated Neural Network can be used in order to prevent the vanishing of the early gradient through the network (116). We chose arbitrarily the Gated Recurrent Unit (GRU) (117) for the RNN model. An alternative strategy could consist in using a Long-Short Term Memory (LSTM) (118), and we leave the comparison between these various GNN for future studies. For the sake of sparsity and because it is not relevant to the present study, we do not detail these structures here, and refer to the corresponding papers for further details. We chose a hyperbolic tangent function activation function for all these networks, which proved to be of greater accuracy than the Restricted Linear Unit (119; 120) and softplus activators.

The RAE-DDM was configured with 50 nodes per layer in each perceptron, GRU included, for both the *i.i.d*.-VB and RAE-DDM models. For the latter, the length of the latent variable **z***_j,n_* was set to 4 for all experiments (so was the length of the subject-specific latent variables **ζ***_n_*), as it showed to provide a sufficient predictive accuracy during piloting. A four component (or layer) normalizing flow was used for the *i.i.d.-*VB and RAE-DDM approximate posteriors.

Each of the VB models were coded in Julia (121), optimized with a noise-insensitive version of the Adam optimizer (58; 122) for 80000 iterations, in parallel with an implementation of the SGD algorithm similar to (123), over 8 CPUs node of various kinds (IvyBridge, Xeon, E5-2650v2, 2.60GHz; Skylake, Xeon, 4116, 2.10GHz; SandyBridge, Xeon, E5-4620, 2.20GHz) with no accelerating device such as GPUs. The code will be made available online.

The *i.i.d*. versions of the DDM (VB or HMC) were all tested with a model identical to the one used to generate the data, i.e. where the drift was influenced by the perceived motion.

### 3.3. Testing for model specifics: Convergence and overfitting assessment

We tested the convergence of the two *i.i.d*.-VB algorithms (with either fixed or variable thresholds) using a validation dataset during training. Convergence rate was fast, and the performance over the validation dataset worsened slightly during the optimization up to a certain plateau. Note that, in piloting experiments, we found that this effect was mitigated by the use of Variational Dropout. This effect was not observed for the *i.i.d*.-VB models.

We calculated *a posteriori* an ad-hoc measure of VB performance named Pareto Smoothed Importance Sampling (PSIS) (124). PSIS provides a parameter 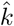 that gives an estimate of the goodness of fit of VB once the optimization procedure has been performed, and also provides a way to refine the accuracy of the posterior estimates (we refer to (125) for a full description of the method). In short, if 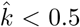, one can infer that VB achieved a good performance for the task. Both *i.i.d*.-VB models showed a very good performance, with a maximum value of 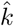 of 0.51 for the classical versions of *i.i.d*.-VB and 0.41 for the full versions. Figure 7 compares the convergence of the three VB methods.

**Figure 5:**
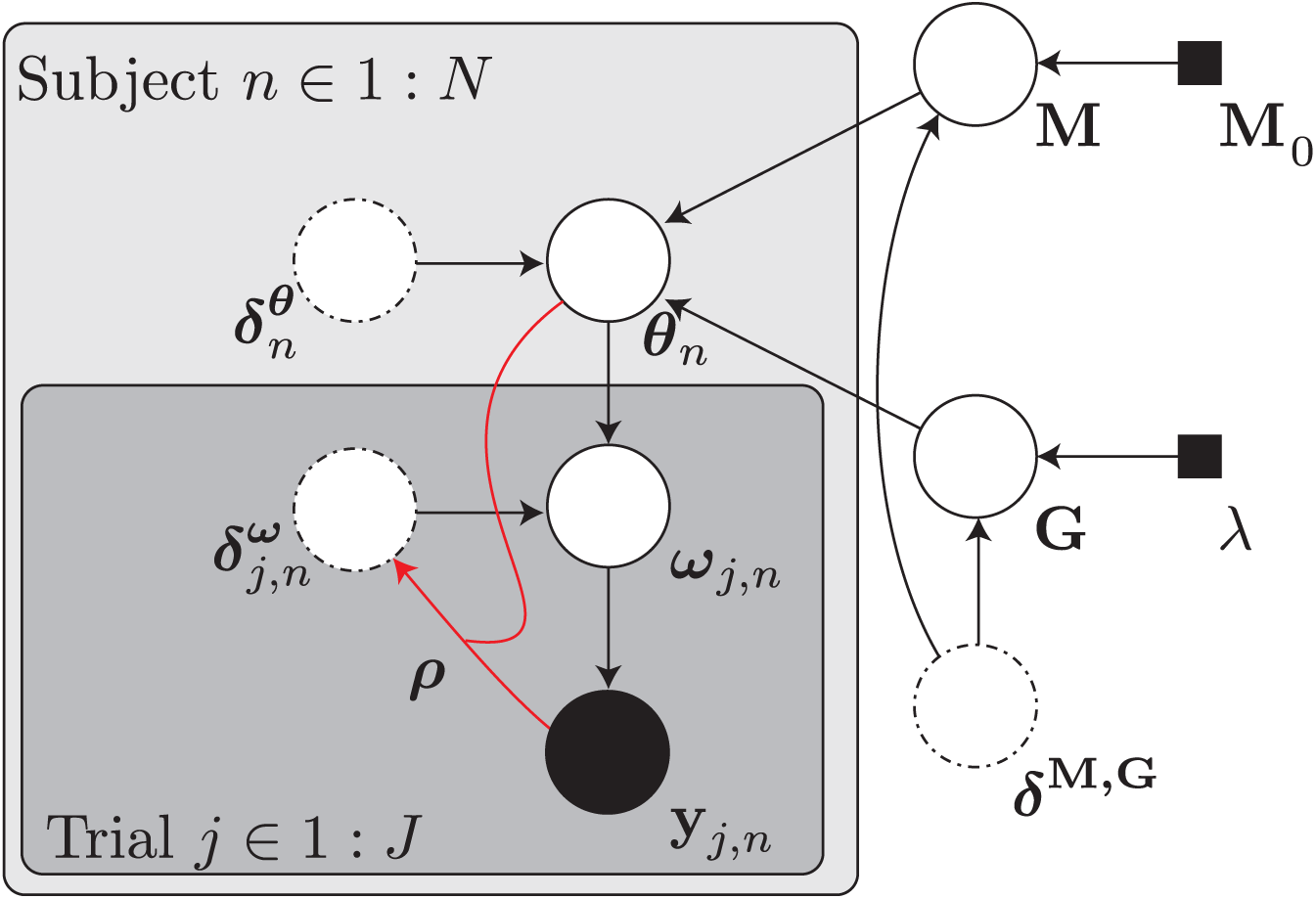
*i.i.d*.-VB and HMC full hierarchical structure. The DDM parameters were distributed according to a subject-specific prior 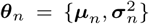. In turn, these parameters were distributed according to a latent Normal M prior 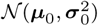 and a latent Inverse-Gamma **G** prior 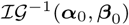 distributions, respectively. The hyperprior of the first distribution was chosen to be a Normal Inverse-Gamma distribution M_0_ with parameters 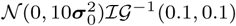. The hyperprior of the second distribution was a set of Exponential distributions with shape parameters λ = 0.01. These values were chosen in order to be weakly informative. This model was identical for the VB and HMC tests. However, in the VB implementation, the approximate posteriors were modelled according to a 4-layers Normalizing Flow with parameters ***δ***ֹ, where · indicates the latent variable. The parameters 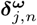 were generated thanks to an inference network with parameters *ρ*.

**Figure 6:**
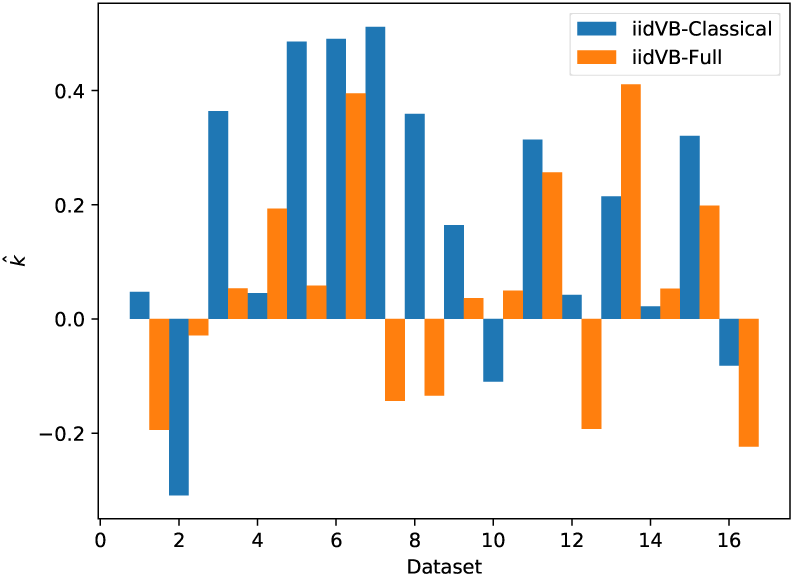
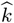 value for each dataset.

**Figure 7:**
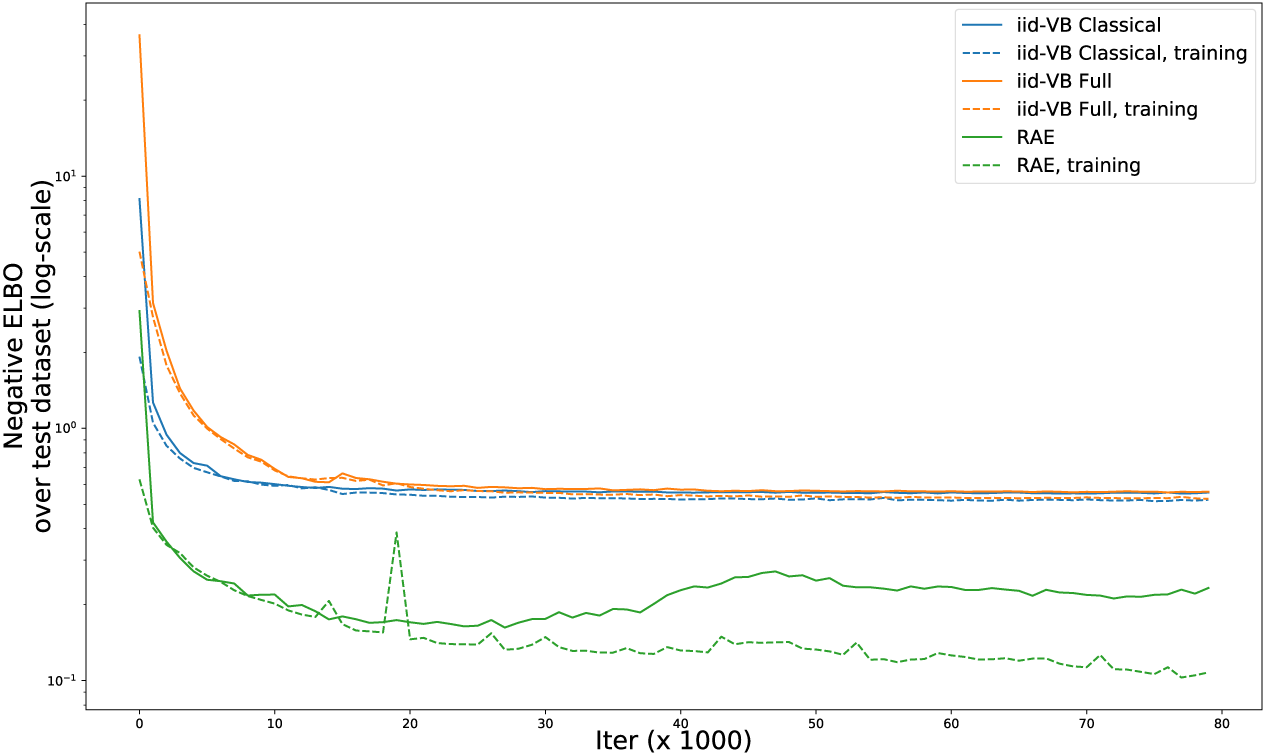
Convergence rate on a validation dataset (plain) and training dataset (dashed) for the *i.i.d*.-VB and RAE-DDM approaches.

**Figure 8:**
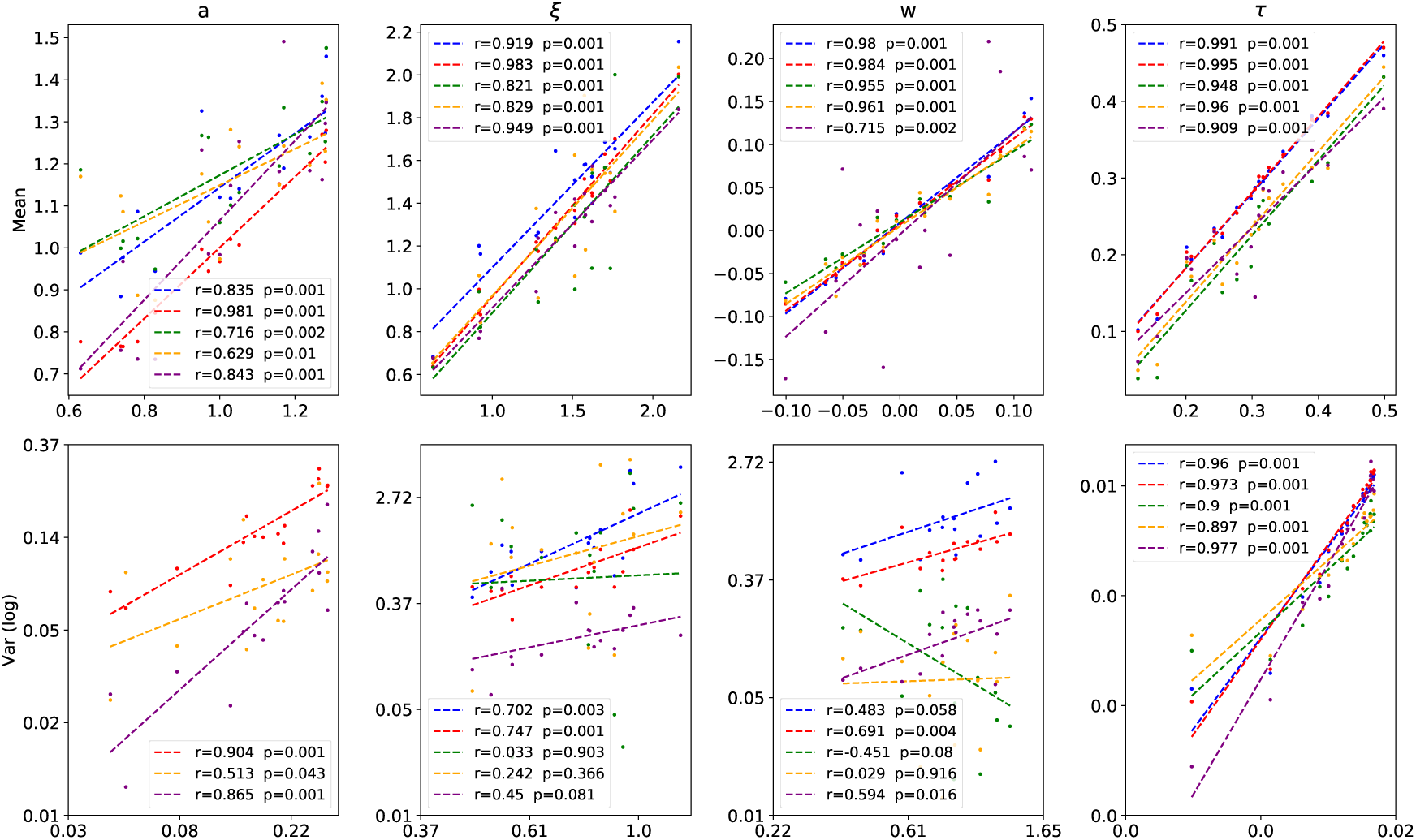
Population value correlations. **Blue**: HMC-Classical, **Red**: HMC-Full, **Green**: *i.i.d*.-VB -Classical, Yellow: *i.i.d*.-VB -Full, **Purple**: RAE-DDM. Fit of the Classical versions of HMC and *i.i.d*.-VB are lacking in the lower left panel, because these models did not account for threshold variability.

To compare the three VB models, we ran a vanilla Bayesian Model selection algorithm ((72)) to investigate the comparative probability that each model generated the data. This method assigns individuals (datasets in this case) to competing models, and can provide an exceedance probability *ϕ* which can roughly be interpreted as the probability that a model is better than all others. This analysis clearly favored the RAE-DDM with a total of 13.27 datasets (*ϕ*=0.999) assigned to this model, with respect to 1.39 and 1.34 datasets for the classical and full versions of the *i.i.d*.-VB model, respectively. Interestingly, these results did not favour any of the two *i.i.d*.-VB models.

### 3.4. Fit Comparison

We tested the fit of the five models on the training dataset, as HMC does not provide the opportunity to generalize the inference to unseen datapoints. We report here the results in terms of Pearson’s correlation coefficients. Euclidean distance (or log-Euclidean distance for the variances) is reported in the Appendix Appendix D. All results are reported in an unbounded space (a, w are mapped with the inverse mapping described in Appendix A, and all statistics involving the variances were computed with the log-value of the variance). For the RAE-DDM, means and variances were computed according to Equation (B.6). Finally, all the results that regard the drift rate were corrected for motion direction.

*Population-wise fit*. At the population level, HMC performed well for all parameters average values wrt VB-based methods. From a correlation point of view, the RAE-DDM had an acceptable performance, although it tended to underestimate the variance of each parameter.

Quite interestingly, the HMC-Full performed well in discriminating datasets with high and low variance of the *a* and *w*, which would not have been possible if the variance of these two parameters were equivalent, as claimed in earlier studies ((7)). Moreover, fit of the HMC-Full was always equal or better than the classical version with no threshold variability, which confirms that the Full-DDM with threshold variability is identifiable at the population level.

*Subject-wise fit*. All methods provided subject-specific estimates of the mean parameters that correlated well with their true value, with the notable exception of *τ*, whose fit was acceptable only for the RAE-DDM. In all mean parameter estimations, the RAE-DDM outperformed or was undifferentiable (for *w*) from other methods.

Even more important is the difference between the evaluation of the subjects’ specific trial-to-trial variability of the four DDM parameters by the five models. First, it can be clearly seen in Figure 9 that the RAE was able to assess variability with a reasonably high accuracy that was inaccessible to other methods. Second, the models (HMC or *i.i.d*.-VB) where the threshold variability were implemented did *not* perform significantly worse than others: on the contrary, the HMC-Full uncovered better the variability of the drift, start point and non-decision time than his classical counterpart. These observations suggest again that relaxing the *i.i.d*. assumption and including the threshold variability in the DDM might be beneficial.

**Figure 9:**
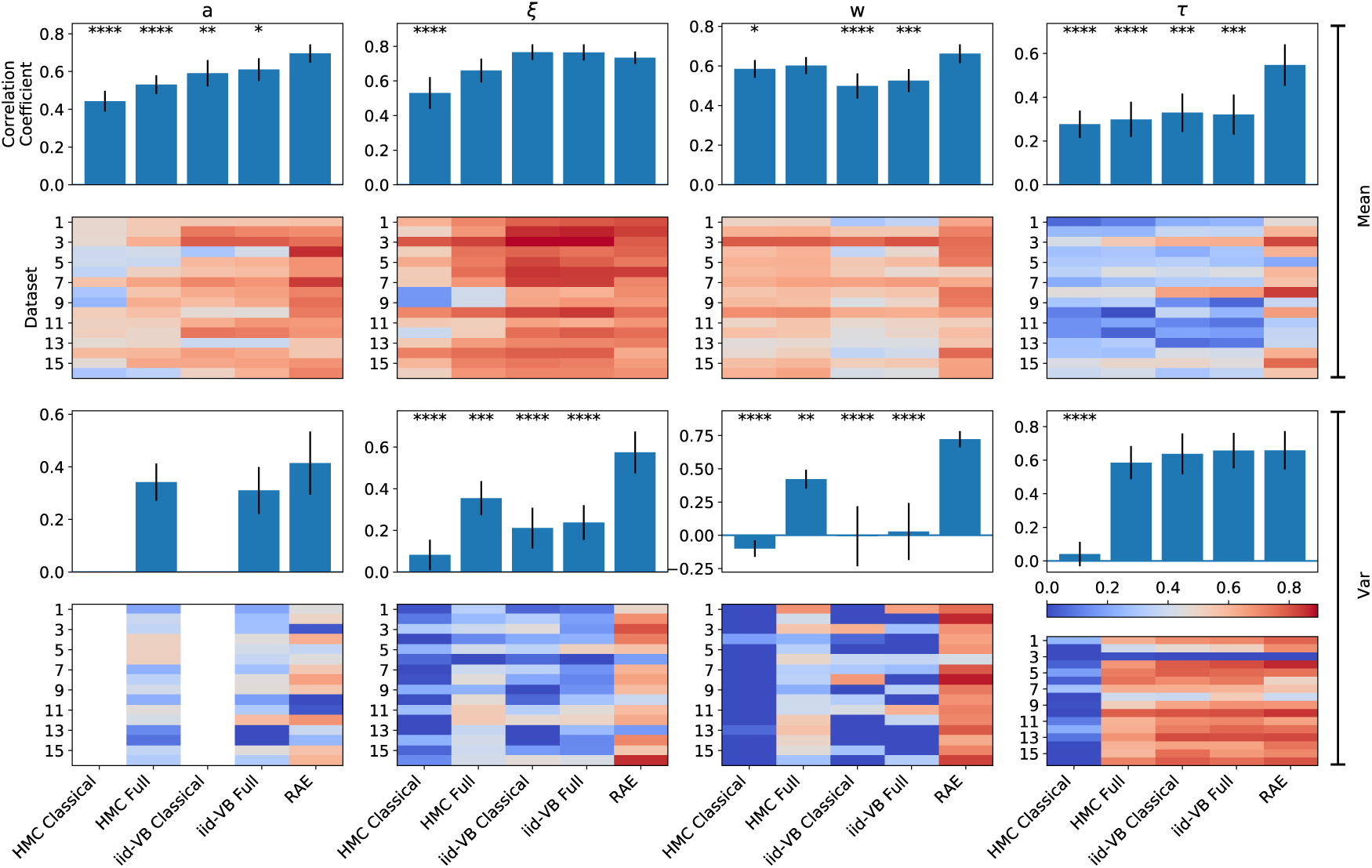
Subject value correlations. Heat plots show the value of the Pearson correlation coefficient for each dataset, for each model. The bar plots on top of these show the average of the corresponding heat plots. Error bars show the value of one standard deviation. The stars indicate the p-value range of a GLM model that accounted for the value of the true-to-estimate correlation coefficient value as a function of the fitting method (* for *p* < 0.5, ** for *p* < 0.01, *** for *p* < 0.001 and **** for *p* < 0.0001). It can be observed that when the fit of the RAE-DDM was poor, it was usually also the case for other methods.

*i.i.d*.-VB methods had reasonable performance for mean estimates but were poor at estimating trial-wise variability of the DDM parameters.

*Trial-wise fit*. The RAE-DDM performed better at the trial level than each and every other model (Figure 10). Overall, *i.i.d*.-VB and HMC performed equally well for all parameters. The correlation at the trial level of the parameter *a* was slightly lower than others. Whether this difference was due to a lower relative variability of this parameter wrt. the others, or to the assumptions about the data-generation process remains to be explored.

**Figure 10:**
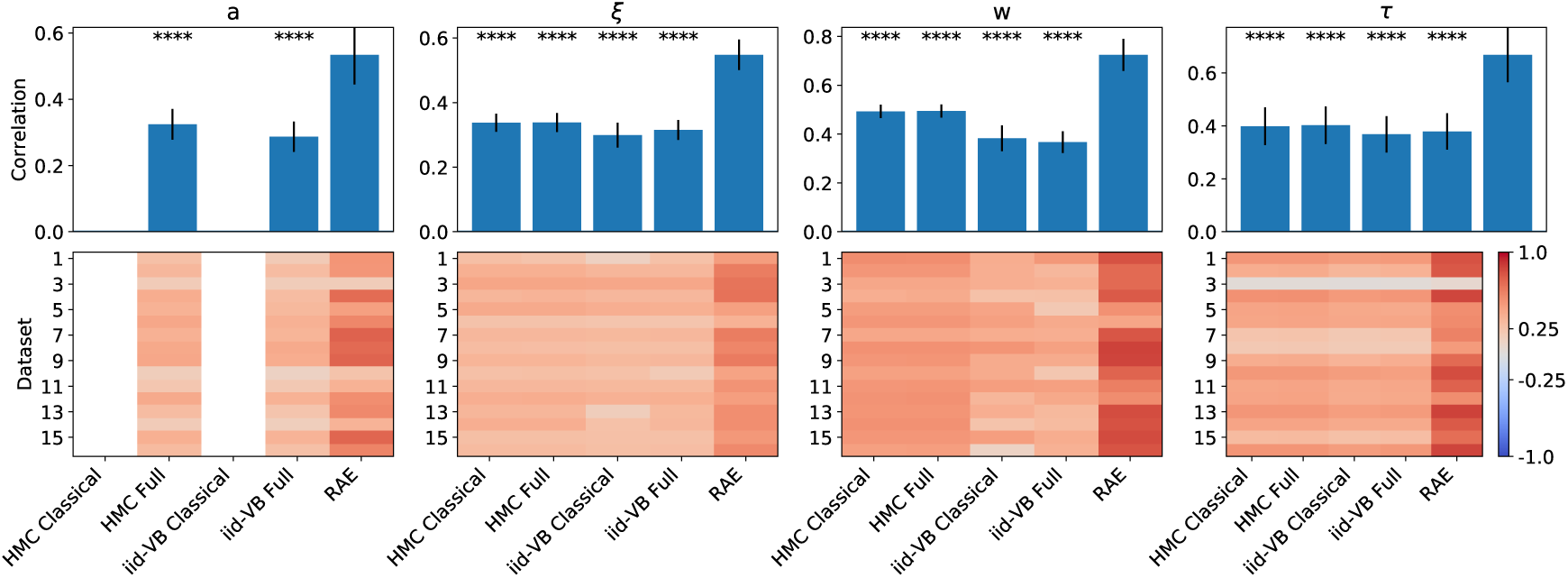
Trial-value correlations (averaged over subjects for each dataset).

Nevertheless, the RAE-DDM was able to assess the value of the *a* at each trial with a relatively good accuracy (Figure 11).

**Figure 11:**
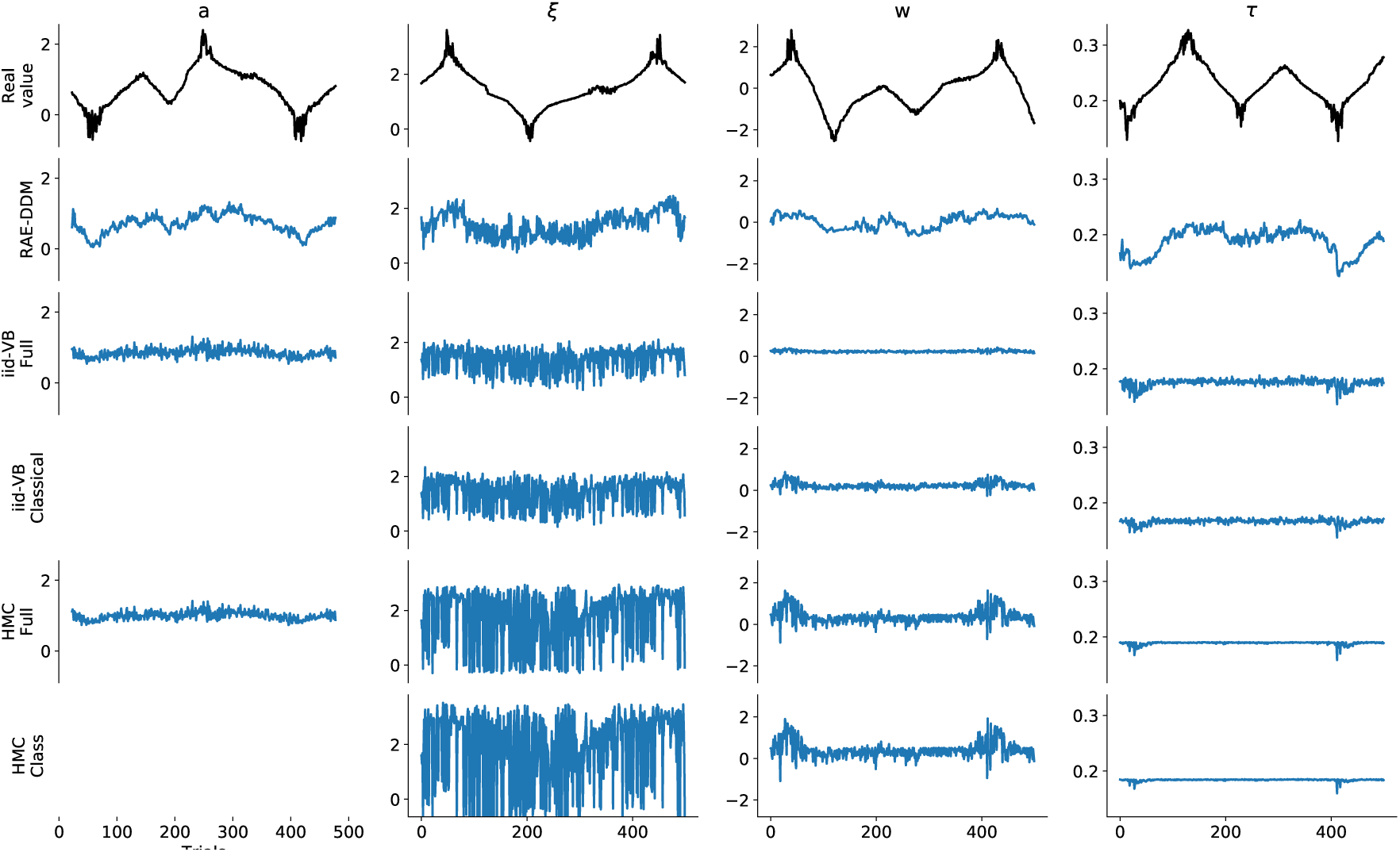
Randomly selected subject parameter values, and fit for each model. The RAE-DDM estimated the parameter trace with a high accuracy. We emphasize that the fit of the drift (corrected for motion direction in the figure) followed the motion direction pattern that was used to generate the data with a high accuracy, even though this was not directly instructed in the model. Also, other parameters did not follow this pattern, meaning that the algorithm was able to discriminate which parameter value did and did not depend upon the motion direction without explicit user input. This contrasts with the four other methods, where the effect of motion direction was explicitly implemented in the model. It is noteworthy that beside their lack of capacity to account for the pattern of variation of the parameters, the four other methods also failed to provide a good estimate of the parameter variance in this example.

## 4. Discussion

In this paper, we present an advanced, scalable, data-driven algorithm to fit the Drift Diffusion Model (DDM) in a fully Bayesian setting that can recover the parameter value at the trial, subject and population levels. This algorithm, named the Recurrent Auto-encoding DDM (RAE-DDM) mainly relies on recent advances in Variational Bayes Inference (VB). The main assumption of this model is that the data are correlated in a forward manner, i.e. that the current value of the data and parameters are conditioned on the preceding value of the data and parameters. It relaxes the *i.i.d*. assumption about the parameter distribution, as well as the assumptions about the distribution of these parameters and, most importantly, about the stationarity of the threshold parameter. We show that the fit of this model outperforms other methods (*i.i.d*.-VB and Hamiltonian Monte Carlo (HMC)), and that inference can be efficiently achieved at every level of the hierarchy with correlation coefficients and distances between true and estimated values that are impossible to obtain with previous approaches.

The RAE-DDM enjoys countless advantages; it can be optimized in parallel or on GPU, it requires a low memory usage thanks to the use of Amortized Variational Inference, and, for the same reason, it is generalizable to unseen data. Its structure allows it to be optimized in an online manner. Furthermore, by the use of dimentionality reduction and variational dropout, this model prevents overfitting.

It can account for the effect of regressors of any kind, without requiring to precisely set the relation that ties these regressors to the parameters. Yet, it provides useful information about the parameter values: since the quality of the fit can be expected to be high for a wide range of datasets, the precise estimation of the trial-wise parameters value allows the user to adequately interpret the parameters fit *a posteriori*, based on the trial-to-trial changes in input regressors. This contrasts with classical, non data-driven modeling approaches, where the model structure has to be set beforehand, fitted and then interpreted based on the quality of fit. Therefore, the RAE-DDM comprimises informativeness of the model with assumptions about the model specifications.

Importantly, we do not claim that informative models should not be used. We suggest that, if an informative model needs to be tested, which will still be desirable in most instances, the universality of the RAE-DDM can provide a benchmark of the parameters fit and an upper bound to the model evidence that can be compared with the informative model fit, with closer fits between the two approaches providing empirical evidence in favour of the grounds of the informative model.

It is noteworthy that, contrary to previous approaches (5; 12; 27; 7; 29; 30; 48; 16), here we did not omit any parameter when modelling the trial-by-trial variability. An important contribution of the present paper is indeed to show that the threshold variability can, and hence should, be included in the model trial-to-trial autocorrelations are included in the model. The present method is the first, to our knowledge, in which this specific feature, previously regarded as unidentifiable, has been implemented. Despite the inclusion of this feature in our algorithm, we observed a better fit of the whole set of parameters, showing that its addition did not compromise, but actually improved the identifiability of our model.

Also, while testing the inclusion of the threshold in the models in several datasets fitted with the *i.i.d*.-VB and MCMC approach, we found that not only the RAE-DDM was able to uncover the value of the threshold variability, but that the versions of the HMC or *i.i.d*.-VB models that accounted for threshold variability did not perform worse than others in recovering the parameters of the model. This simple observation should motivate researchers to take with more caution the *a-priori* statement that the threshold variability should be discarded. However, we acknowledge the fact that the exact similarity between the true generative model and the fitted model specifications may have biased these results: as it has been shown by (23), the choice of a different prior distribution for *a* and *w* would probably have lead to different, less accurate results. Nevertheless, Bayesian Model Selection and cross-validation should be used to assess the quality of the fit.

The first innovation of our models is the use of VB to solve the Full DDM problem. Recent advances in VB have made it nearly as precise as MCMC (103; 64; 126; 80), easy to use (127; 98; 88) and adapted to potentially very large problems (76; 70; 65). We used these advantages of VB to solve two challenging problems: the first one being to recover the DDM parameter variability from trial to trial (7) with a fast algorithm; the second being to recover an estimate of the trial-to-trial cross-correlation of the DDM parameters and related behavioural data.

Another key feature of the RAE-DDM is its ability to make inference at the trial level. The applications of this feature are countless. For instance, one could assess how parameters change over time following pharmacological or neurophysiological intervention (e.g. dopamine modulation (128) or sub-thalamic nucleus stimulation (40)). It also makes it possible to determine the relationship between neurophysiological data (EEG, fMRI) and trial-by-trial values of DDM parameters, similarly to previous studies in which the post-error slowing phenomenon was related to Frontal Theta bands and the decision threshold was shown to correlate with this signal (129; 40; 130). The present methods open the door to a more precise study of such phenomena, by allowing precise estimation of the generative parameters in a data-driven manner.

The highly generic structure of the RAE-DDM opens the door to many further developments. A potential improvement of the present approaches would be to broaden the models to other sequential sampling models. Many of them have a closed-form FPT density function formula (131), allowing our models to extend to these other frameworks. Applicable models include the Collapsing Barrier DDM (132; 133) or the Time-varying Ornstein-Uhlenbeck model (134; 135; 1; 136).

Also, many appealing features could be implemented with the RAE-DDM to model neurophysiological measures together with behavioural data. Similarly to (55), we could set the physiological measure (e.g. ERP, pupil measure, fMRI measure) to be the output of the generative model.

Finally, the model we propose is scalable at the trial level, but the subject-specific latent variables **ζ***_n_* approximate posterior still needs to be fitted independently. Scalable inference at the subject level would probably greatly improve the performance and efficiency of the model.

The assumptions of the RAE-DDM should be discussed here. The first and most obvious assumption is that the parameters and the behavioural results are cross-correlated between trials in a forward manner. As discussed above, many behavioural observations are known to reflect complex psychological biases, which justifies the use of previous parameter values as regressors for the current one. A classical way to test which parameter of the DDM is impacted by the previous behaviour would be to explicitly model the supposed cross-dependency among parameters (40). Thanks to its generic and flexible form, the RAE-DDM automatically selects the model configuration that explains the most variance up to a predefined degree of complexity (represented by the length of the latent variable **z***_j,n_*). The results we provide show that the presence of this cross-correlation does not harm the fit, but constrain it adequately, as only very few parameter sequences will explain a specific behavioural pattern across the whole experiment.

The second assumption of the model is that the parameters covary at the trial level in a complex way. In the RAE-DDM, this can be observed by the fact that the latent variable **z***_j,n_* is mapped onto ***ω****_j,n_* with a complex non-linear function. This, again, contrasts with most approaches where the variance of the DDM parameters is assumed to be independent. We think that this latter assumption is likely to be incorrect: for instance, after an error, a subject will likely modify both his bias and threshold value, and therefore these two parameters variability should not reasonably be considered as independent. Once more, the richness of the method we propose does improve rather than impair the fit quality by constraining parameter values to lie within a certain correlated subspace.

In summary, we present a VB-backed hierarchical, sequential and data-driven implementation of the DDM that can retrieve relevant information of the data-generating process at each level. We focused our attention on the possibility of modelling the trial-to-trial dependency of parameters through the development of Recurrent Auto-Encoders. We show that this model is highly flexible, scalable and much more precise than other classical approaches provided that there exists some form of auto-correlation in the values of the parameters and/or data.

## Acknowledgement

This work was performed at the Institute of Neuroscience (IoNS) of the Université catholique de Louvain (Brussels, Belgium); it was supported by grants from the ARC (Actions de Recherche Concerté es, Communauté Francaise de Belgique), from the Fondation Médicale Reine Elisabeth (FMRE), and from the Fonds de la Recherche Scientifique (F.R.S.-FNRS) to A.Z. A.Z. was a Senior Research Associate supported by INNOVIRIS and is currently 1st grade researcher for the french National Center for Scientific Research (CNRS).

This research used resources of the Center for High Performance Computing and Mass Storage (CISM, https://uclouvain.be/en/research/cism) located at the Université Catholique de Louvain, Belgium, which is supported by the Fonds de la Recherche Scientifique – Fonds National de la Recherche Scientifique (F.R.S.-FNRS). The CISM is member of the ”Consortium des Équipements de Calcul Intensif (CÉCI)” (http://www.ceci-hpc.be).

## Appendix A. Normal prior for the DDM parameters

If we are to put a normal prior on the DDM parameters, it is important to transform the threshold, starting point and non-decision time such that they lie in an unbounded space. Consider the set of bounded parameters {*a,ξ,w,τ*} where 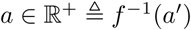, 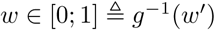, 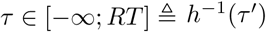 and 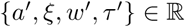. We can construct a distribution

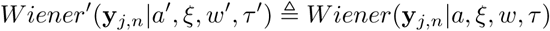

that satisfies this condition for *a, w* and *τ* by choosing appropriate transforms *f*, *g* and *h*. A natural choice for *f* and *g* are respectively the exponential or softplus function – the latter being more advantageous since its gradient is bounded to one, which avoids numerical overflow for samples with large values of the threshold – and some sigmoid function such that *σ*(*wʹ*): ℝ → [0; 1].

## Appendix B. Full DDM as an Empirical Bayes problem

If we are to use the MLE method for the Full DDM for a set of *N* subjects with *J* trials per subject, we therefore need to solve the following optimization problem:

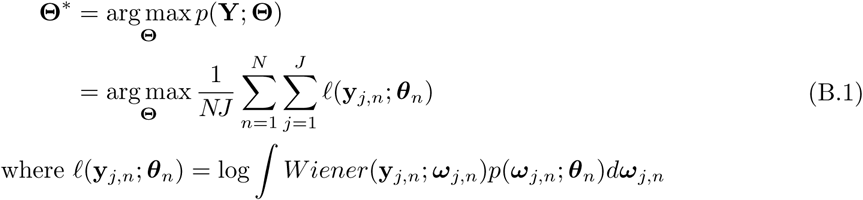

For simplicity, we will consider that the DDM parameters all lie in an unbounded space, so that their prior probability can be formulated as a Gaussian with parameters 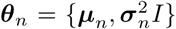, with *I* being the identity matrix (see Appendix A for justification and implementation).

In Equation (B.1), we have used the average log-likelihood with the purpose of showing that, for a sufficiently large number of samples *NJ*, this average log-likelihood takes the form of a sample of the true expected log-likelihood function:

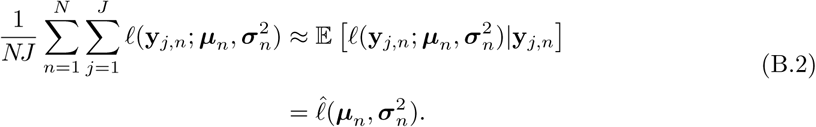

Since the integral in Equation (B.1) has no analytical form for *a, w* and *τ*, and since we need to evaluate it for every trial (12), finding the MLE of 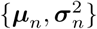 will require us to compute the gradient of each trial using numerical integration over three dimensions (or two in the classical version of the Full DDM), which we can expect to be computationally expensive.

We now show how Bayesian Statistics arise as a natural way to deal with this problem. The maxima of the likelihood function are located where the value of the gradient of the expression in Equation (B.2) is equal to 0:

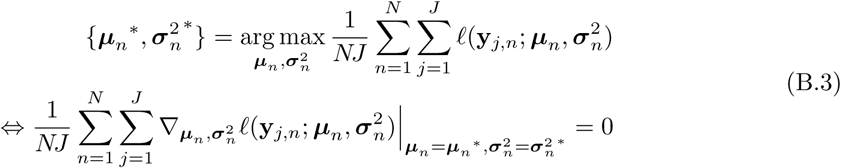

We can unpack this expression for each trial and make use of the Fisher’s identity (137) to show that:

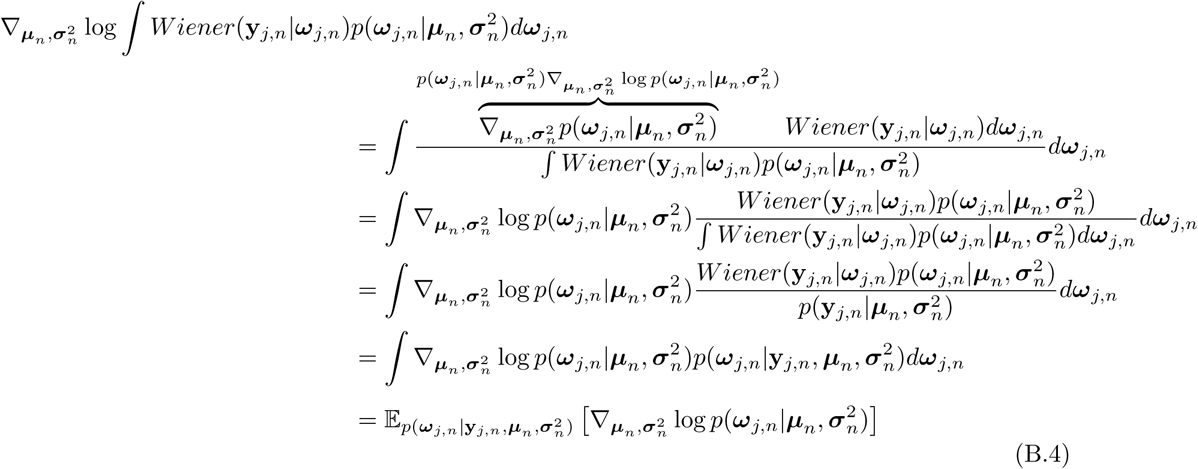

where we have used the Leibniz rule, the log-derivative trick and finally the Bayes theorem 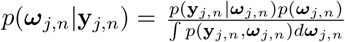. Equation (B.4) shows that the DDM problem can be solved if we can compute (or approximate) the posterior probability 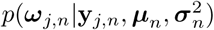. Indeed, since the gradient formulated in Equation (B.4) must be zero at the mode, we can expand this formula and apply it to all trials to show the following:

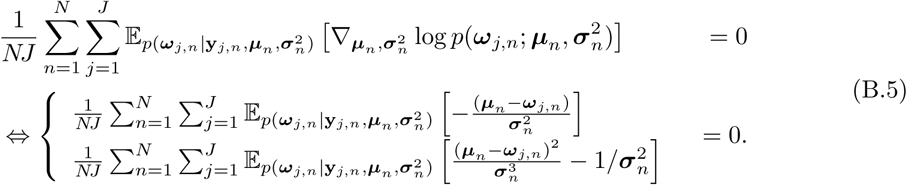

Solving for *μ_n_* and 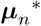 we have:

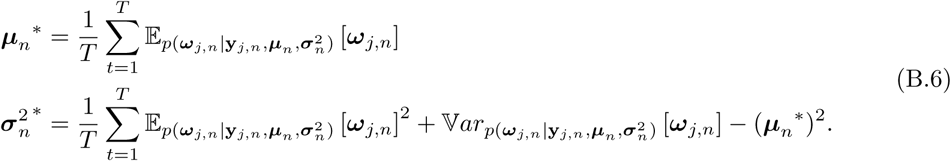

This can be viewed as an Empirical Bayes problem (138; 139) where a MLE of the prior distribution 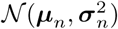 has to be retrieved from the data. Using a Hierarchical Bayesian terminology, ***ω****_j,n_* is the latent (or local) variable and 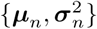 are the prior or global parameters (Figure 2).

Equation (B.6) shows that 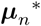 and 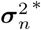 will have a closed form formula as long as we can estimate the first and second moments of 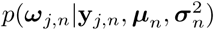 for each trial. If the posterior probability of ***ω****_j,n_* were easy to estimate, this cross-dependency between the posterior probability over ***ω****_j,n_* and the value of 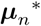 and 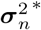 could suggests an expectation maximization (EM) algorithm: we could iteratively estimate the posterior 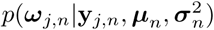 under the current belief of *μ_n_* and 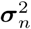, and then, we could estimate 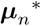 and 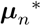 using Equation (B.6), until some convergence criterion is reached. This posterior probability does not, however, have a closed-form: additional work is therefore necessary to infer the value of the posterior probability.

## Appendix C. Approximate posterior Specification

### Appendix C.1. Approximate posterior shape and reparametrization trick

Sampling the first three parameters of the DDM is trivial if these are distributed according to a (possibly diagonal) Gaussian distribution. Let 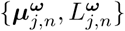 be the mean and lower-Cholesky decomposition of the amortized approximate posterior Gaussian distribution *q*(***ω****_j,n_*|**y***_j,n_*, ***θ***_n_; *χ*), where 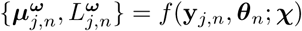, then:

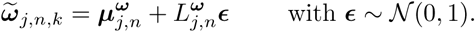

One can easily see that this expression is differentiable and therefore suited for SGVB or IWAE.

Sampling the non-decision time is more tricky: whereas the domain of the prior is unbounded (τ*_j,n_* ∈ [−∞; ∞]), the domain of the approximate posterior we would wish to derive is not (τ*_j,n_* ∈ [−∞; *t_t_*]), as an unbounded probability distribution would generate samples that might give a gradient of 0 to the ELBO, which would in these case take a value of −∞.

One solution to this problem would be to use a Truncated Gaussian 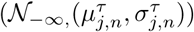 distribution, where the upper bound would be set to the RT of interest, and the lower bound would be set to −∞. Two main reasons might discourage us from adopting this strategy: the first would be that efficient random number generation algorithms for the truncated Gaussian are not differentiable, because they rely on acceptance-rejection algorithms (106). However, as the cumulative density function (CDF) of the univariate truncated Gaussian can be computed efficiently, we can compute an estimator of the gradient 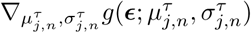 as (108):

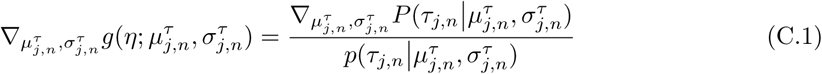

where *P* is the CDF of 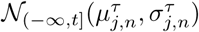 and *p* the PDF. We refer to the code available online for the derivation of this gradient. Note that, whereas for the multivariate Gaussian, the gradient and random numbers were computed at the same time, here, one must first generate the random number and then, on this basis, compute the gradient estimator of *g* given the sample that has been drawn.

The second issue why not to use a univariate truncated Gaussian distribution for τ might be found in the strong mean-field assumption that it requires. Using a multivariate truncated Gaussian is, however, much more challenging that factorizing *q*(*τ*) from the other parameters of the DDM, as the CDF of this distribution is extremely hard to derive and expensive compute. Moreover, current implementations of this function are not differentiable wrt the distribution parameters (see for instance (140)). We will introduce how this issue can be solve by the use of Normalizing Flows short after.

### Appendix C.2. Normalizing Flow

If the deterministic mapping from the datapoint to the approximate posterior shape described in Section 2.4.2 facilitates the optimization of the ELBO by reducing the size of the problem, it does not make the approximate posterior more flexible. Here, we used a Normalizing Flow (NF) to transform an initial approximate posterior (typically a Gaussian or uniform distribution) through a sequence of deterministic invertible mappings so that its shape gets closer to the true shape of the posterior (64). The core idea of Normalizing Flows is quite simple: it consists in defining an arbitrarily complex approximate posterior *q*^0^(***ω****_j_,_n_|****δ****_j_,_n_*, χ), where both *δ_j_,_n_* and *χ* are the trial and subject-wise outputs of the inference network.

Consider an initial approximate posterior distribution *q*(*ω*_0_). We would like to transform this probability distribution with a deterministic mapping *ω_K_* = *f* (*ω*_0_, *χ*) with K successive transformations, where *χ* = {χ*_k_* for *k* ∈ 1: *K*} are the parameters controlling how *f* changes the shape of *q*. In order to keep the volume of *q*(*ω*_0_) so that it still integrates to 1 over ***ω****_j,n_* after the sequence of transformations, the resulting probability density function must be multiplied by the absolute value of the determinant of the Jacobian of the inverse transform wrt *ω_K_*, or alternatively divided by the absolute value of the determinant of the transform wrt *ω*_0_:

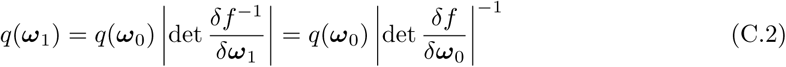

and similarly for the following transformations. After *K* transformations, the ELBO to the model evidence looks like

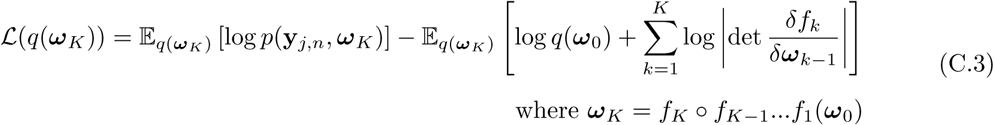

For the sake of simplicity, we chose to use a series of planar transforms with parameters χ*_k_* = {u*_k_*, w*_k_, b_k_*}:

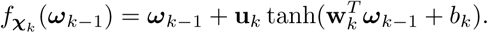

This mapping is only valid if *f_k_*(*ω_k−_*_1_) is invertible. This can be easily achieved if u and w respect specific conditions, which can be found in (64).

*DDM parameters correlation*. The issue of accounting for posterior correlation between *τ* and the other parameters of the DDM finds a convenient solution with the use of planar NF. To account for the covariance between the parameters, we sampled the parameters {*a*, *ξ, w*} according to a diagonal Gaussian distribution, and *τ* according to a truncated Gaussian distribution as stated above. We then transformed the shape of this factorized distribution by a series of planar transformations constituted of expansions and contractions around the initial draw of the parameters 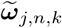. The important point here is that every parameter *except τ* can be transformed in such a way, but always *with τ as an input for the transformation*:

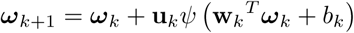

where **u***_k_* and **w***_k_* are two four dimensional vectors, respectively, which are the output of the inference network, with the notable exception that the component of **u**_k_ corresponding to *τ* is equal to 0, which ensures that *τ* lies in the space bounded by the RT. Invertibility of this transformation can still be ensured by the algorithm provided by (64). This form of approximate posterior proved to be efficient in retrieving covariance between *τ* and the other parameters.

### Appendix C.3. FIVO and non-decision-time sampling

The use of the FIVO lead to a convenient sampling algorithm in which we were able to deal with 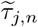 falling above the RT *t_j_,_n_*.

It might however still be the case that each particle of the FIVO was a degenerated sample. To deal with this problem, we artificially modified the gradients of the encoder and decoder parameters by adding a component to the samples of the ELBO such that, if all particles were degenerated at a trial *j*, then the SGD optimizer was instructed to correct the network weights to counter this effect. In practice, the ELBO was transformed to look like:

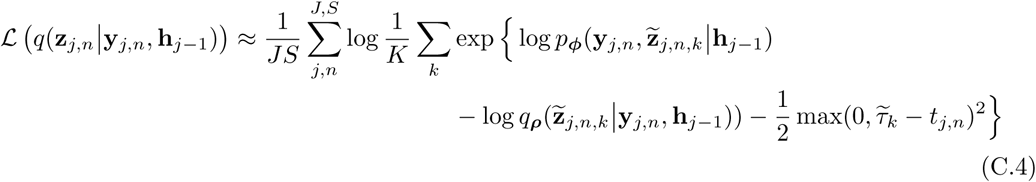

where 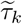 is the sampled non-decision time of the *k* particle. This is a simple case of the use of Lagrange multipliers for constraint optimization: in the case where 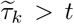 for some but not all particles *k*, then the weight of these particles was set to 0 because 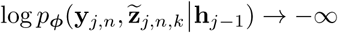. Therefore, the Lagrange correction had no impact on the gradient estimate. The only case where this correction had an impact was therefore when all particles satisfied the above condition, in which case the weights were all set to 1/*K* and the gradient used for backpropagation was set to:

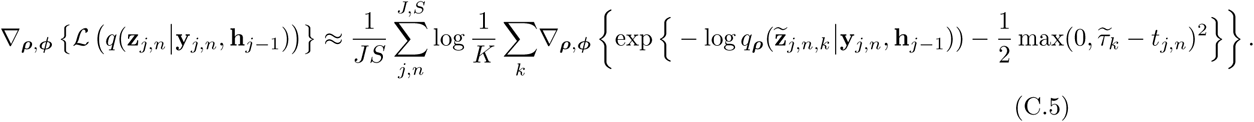

## Appendix D. Euclidean Distances

We report here the Euclidean distances between the true values of the DDM parameters and their model estimates.

Overall, the RAE-DDM performed better or equally than the HMC approach at the trial and subject level.

At the population level (Figure D.12), the mean and variance estimates of the RAE-DDM were slightly more biased than the HMC estimates. This confirms the results shown in Figure 8 that the RAE-DDM had a low discrimination power between datasets.

At the subject level, however, the RAE-DDM performed equally well or better than the HMC Figure D.13. The same was true at the trial level Figure D.14.

We also report here the corresponding correlation plot Figure D.15 of the example subject trace in 11.

**Figure D.12:**
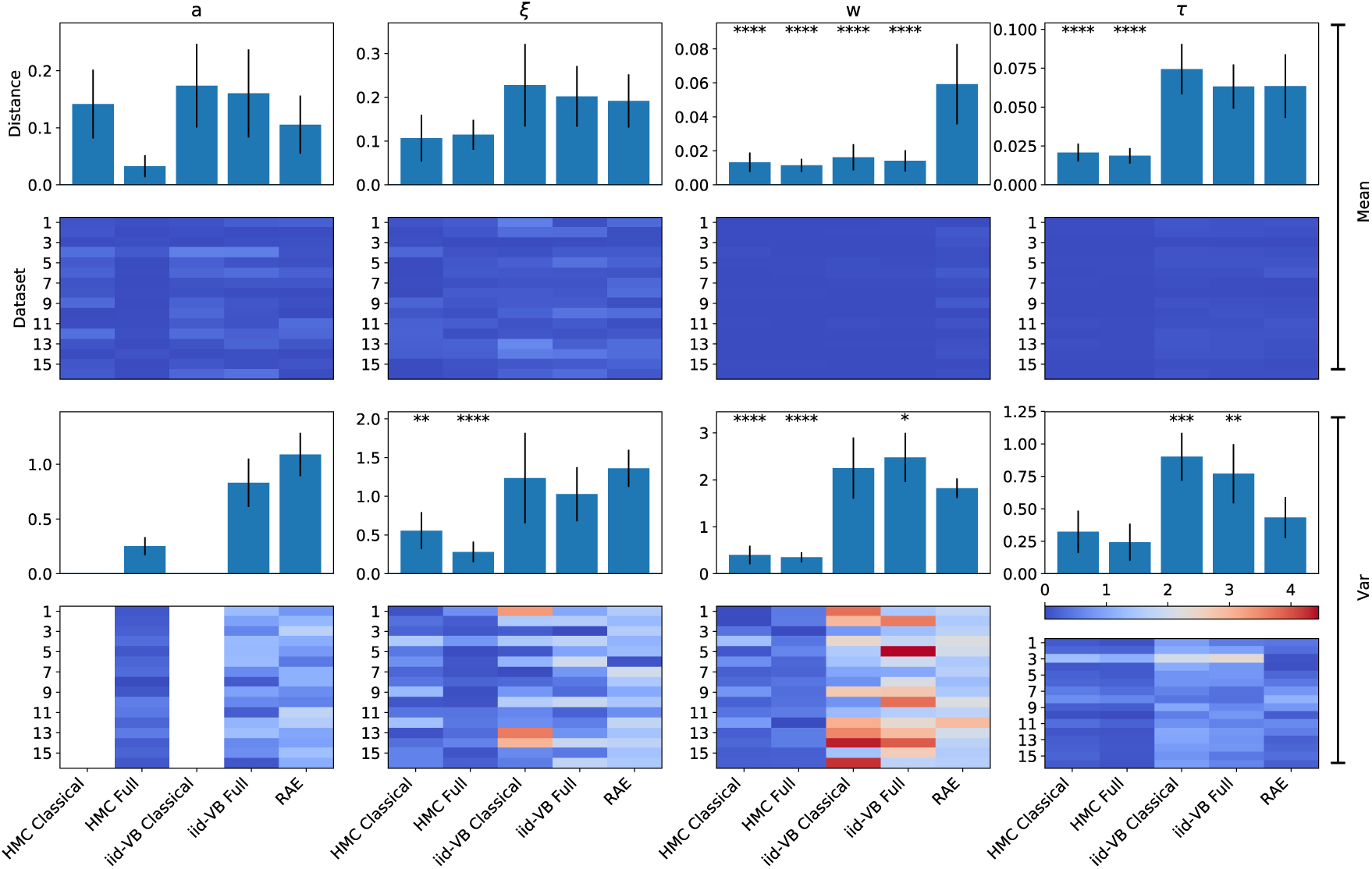
Population true to estimate (log-)Euclidean distances.

**Figure D.13:**
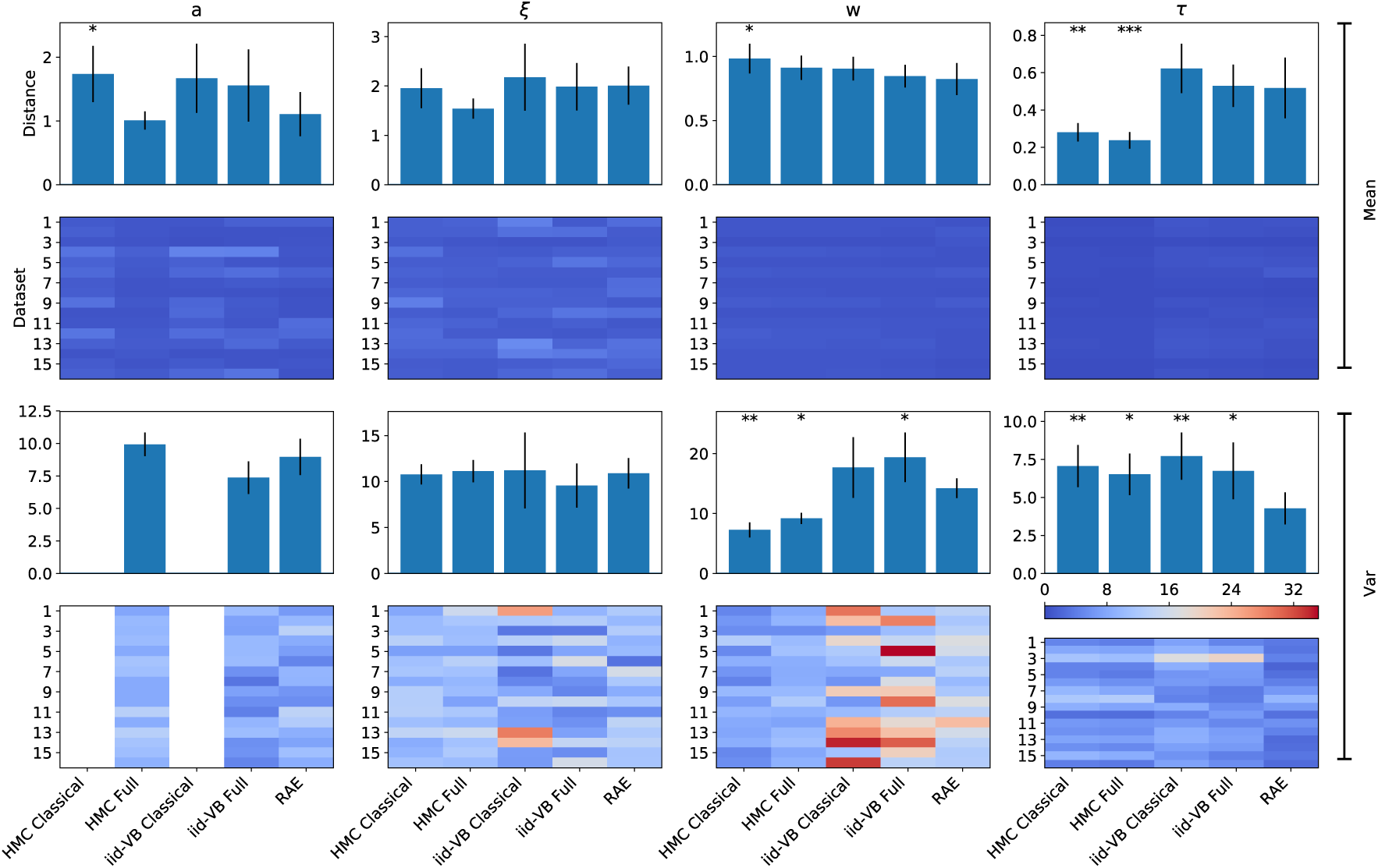
Subject-wise true to estimate (log-)Euclidean sum of distances

**Figure D.14:**
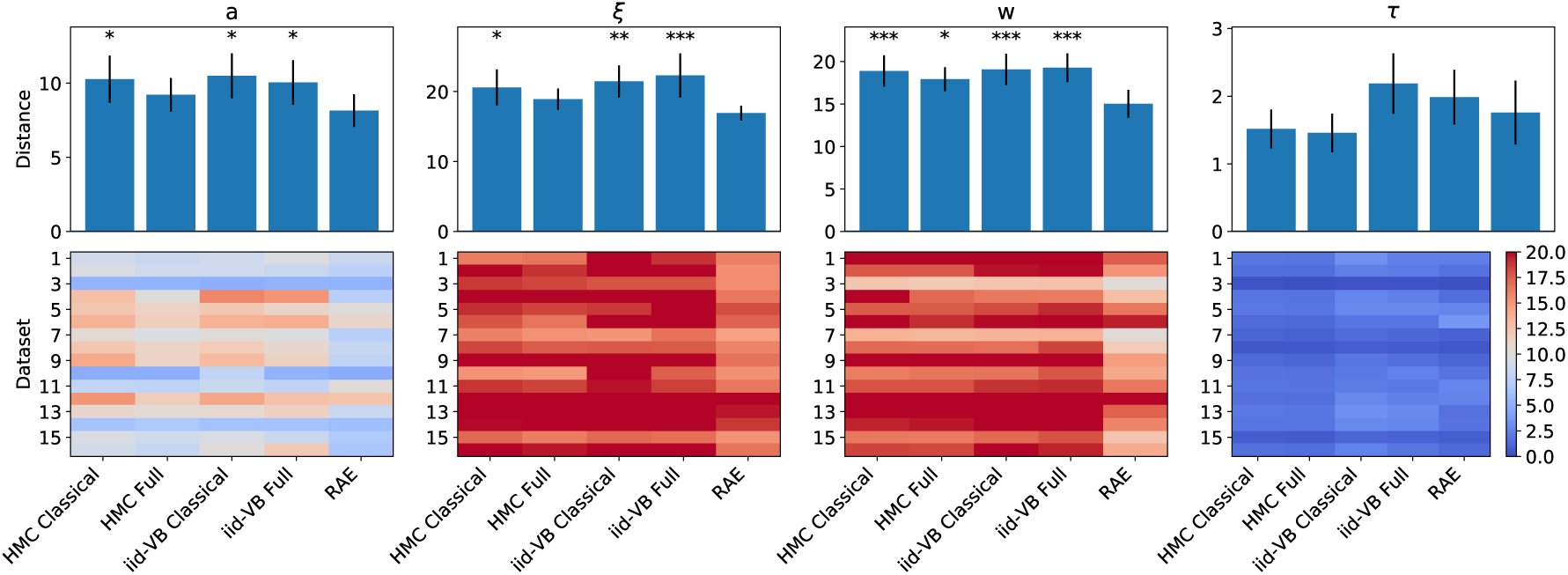
Trial-wise true to estimate (log-)Euclidean sum of distances. The RAE-DDM performed equally well or better than other approaches.

**Figure D.15:**
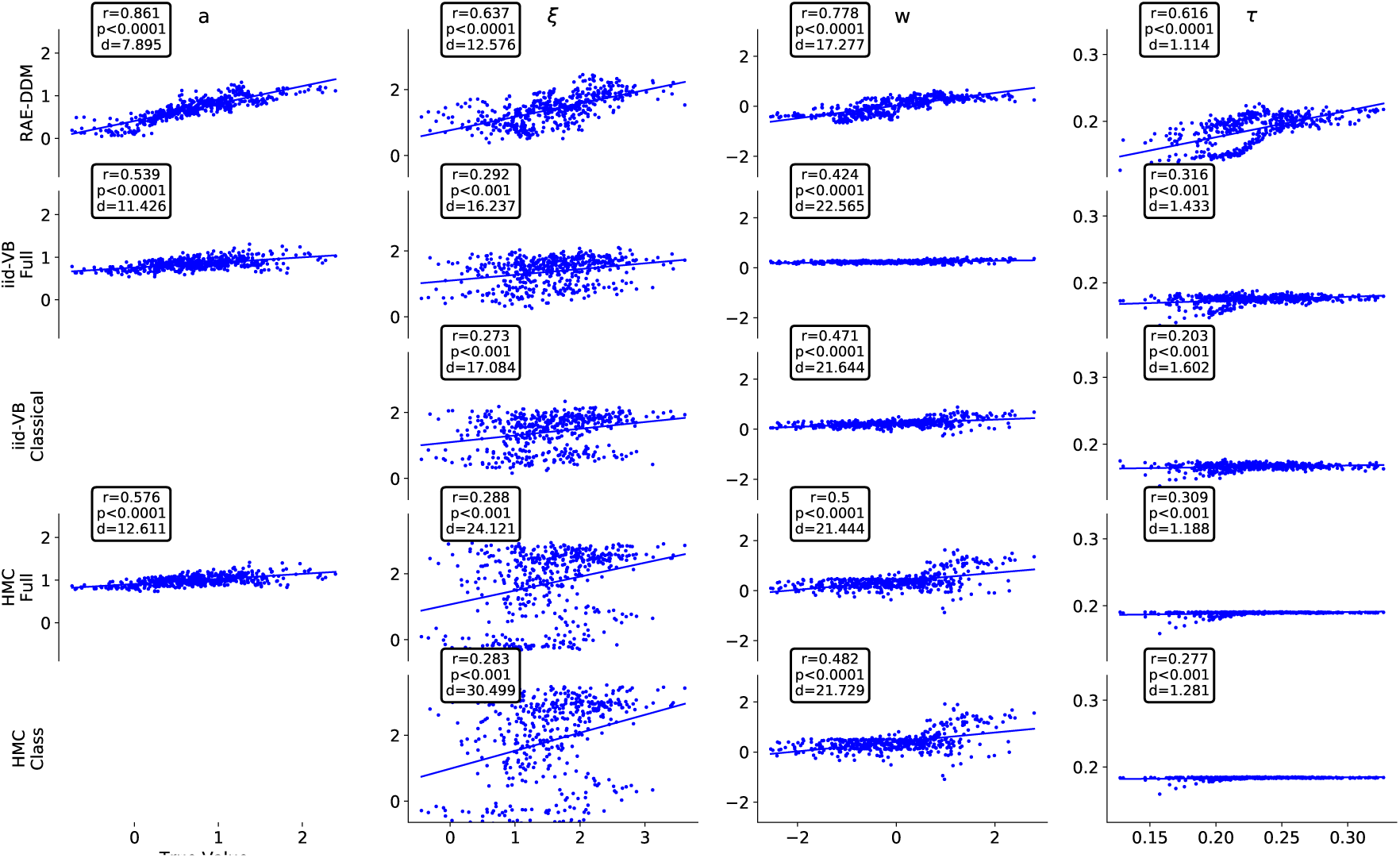
Correlation and distances of the same random subject as in Figure 11. RAE-DDM showed a high accuracy for both these measures, displayed in the upper left corner of each panel. p-values state for the significance of the Pearson’s correlation coefficient displayed above.

